# A fungal pathobiont promotes *Streptococcus agalactiae* vaginal persistence and pathogenesis through physical and metabolic interactions

**DOI:** 10.1101/2025.09.07.674778

**Authors:** Shirli Cohen, Arianne J. Crossen, Melody N. Neely, Kirsty Le Doare, Angela H. Nobbs, Kyla S. Ost, Kelly S. Doran

## Abstract

Complex polymicrobial interactions at the host interface can shape the mucosal landscape and tip the scales between commensalism and pathogenicity. Here, we use a newly adapted murine model of vaginal colonization to show that the human pathobiont *Candida albicans* (Ca) supports Group B *Streptococcus* (GBS) fitness in the vaginal tract and ascension to the uterus. GBS frequently colonizes the vagina asymptomatically; however, during pregnancy, colonization can lead to adverse outcomes and neonatal invasive infection. Using human vaginal isolates of Ca and GBS, we demonstrate that physical interactions contribute to persistence. Triple RNA sequencing of Ca, GBS, and a physiologically relevant model of the human vaginal epithelium reveals that GBS induces arginine biosynthesis in Ca. This drives the expression of bacterial virulence factors and primes GBS for adhesion to the epithelium. We show that interkingdom nutrient exchange can increase GBS pathogenic potential and identify a new target for preventative therapies.

## Introduction

The vaginal tract can host a diverse range of microbes, some of which modulate the commensal or pathogenic lifestyles of one another. Due to their high abundance, extensive research describes the impact of *Lactobacillus* species, both protective and non-protective, on the microbial ecology of the vagina^1–3^. It is less well understood how other microbes influence the landscape, particularly those that are not bacteria, including viruses, archaea, and fungi^4^.

Fungi are highly prevalent members of the vaginal microbial community, colonizing approximately 70% of individuals at least once in their lifetime^5^. Emerging evidence supports that fungi can have extensive impacts on host tolerance of microbes in the vagina as well as on the pathogenic potential of those microbes^6^. The most abundant yeast species in the vaginal tract is *Candida albicans* (Ca)^5,7^, which also colonizes other polymicrobial mucosal sites including the gastrointestinal tract and the oral cavity^8^. Ca vaginal colonization is typically asymptomatic, but a shift in environmental conditions can induce a transition into vulvovaginal candidiasis (VVC), an inflammatory disease accompanied by high neutrophil influx into the vaginal lumen and vulvar pruritus, burning, and pain^9,10^. Ca morphogenesis between budding yeast and filamentous hyphae has been associated with VVC^11^, which may be attributed to the differences in cell wall composition, immune evasive factors, toxin expression, and adhesin repertoires between these two morphotypes. Hyphae express more adhesins than yeast, including several hyphal-specific proteins, which enable them to associate more closely with the epithelial surface and invade through host barriers. Filamentation and adhesin expression are also essential for biofilm formation. These densely packed and highly structured communities can protect Ca from physical and chemical perturbations, antimicrobial drugs, and host immune defenses^12^. Ca adhesins have also been linked to mediating physical interactions with bacteria including *Porphyromonas gingivalis*, *Streptococcus gordonii* and *Staphylococcus aureus*^13–15^. Als3 is among the most well-studied Ca adhesins and drives adherence to both host and bacterial cells among a multitude of other functions^16^.

Like Ca, *Streptococcus agalactiae* (Group B *Streptococcus*; GBS) frequently colonizes the vaginal tract asymptomatically, with a carriage rate of approximately 20% in pregnant individuals^17^. GBS can transition into a more pathogenic lifestyle during pregnancy as it can ascend from the vagina into the uterus and cause a range of adverse pregnancy outcomes including chorioamnionitis, preterm premature rupture of membranes, and stillbirth^18^. Vaginal colonization is also a primary risk factor for transmission to a fetus or neonate, which can result in severe invasive infection^19,20^.

GBS uses numerous adhesive and invasive factors that interact directly with the mucosal surface. This includes pili, PbsP, and BspC, which bind host glycoproteins^21,22^, plasminogen^23^, and multiple components of the vaginal epithelium^24^, respectively.

These factors promote persistence in the female genital tract (FGT), and BspC has additionally been demonstrated to directly contribute to interactions with Ca^25^. Further, GBS encodes more than 20 two-component systems (TCSs) that enable it to sense the environment and regulate complex responses that can tune the GBS lifestyle to fit diverse host niches^26^. Some of these TCSs regulate immune evasive factors such as the capsule, which can help GBS escape recognition and killing by the host. We sought to address whether GBS sensing of and interaction with Ca could impact its lifestyle in the host environment.

In this study, we identify and characterize the molecular relationships between Ca and GBS. Several clinical studies have reported a high frequency of co-occurrence of these organisms^27–32^. These have been sufficient to identify Ca vaginal colonization as a risk factor for GBS carriage. Previous work has also shown that Als3 contributes to interactions with GBS^25^ in co-culture and on monolayers of human vaginal epithelial cells (hVECs); however, the mechanisms governing these interactions are not well understood. Here, we use *in vivo* and *in vitro* models to characterize the physical interactions between Ca and GBS and show how hyphal-bacterial interactions contribute to GBS adhesion to the vaginal epithelium. We use triple RNA sequencing to define unique transcriptomic signatures of Ca and GBS interactions with each other and with the host that result in differential regulation of adhesion and amino acid metabolism. We also demonstrate that Ca promotes GBS vaginal colonization and ascension to the uterus through physical interactions and induction of arginine biosynthesis, identifying arginine as a key regulator of GBS pathogenic potential. This work highlights how interkingdom physical and metabolic interactions can alter microbial ecology, impacting host responses to microbes as well as GBS attachment to and interaction with host tissues.

## Results

### Ca interacts with GBS during vaginal colonization

To investigate the ways that fungi in the vaginal tract can impact the microbial community, we sought to analyze a publicly available dataset that quantifies the abundance of both fungi and bacteria. In a study by Baud et al.^33^, vaginal swabs were collected from pregnant individuals at the time of birth and processed for whole-genome metagenomic sequencing. We accessed this dataset and assessed correlations between fungal and bacterial colonization that were not identified in the report by Baud et al. Those who were positive for Ca were two times more likely to also be positive for GBS compared to the Ca negative population (**Fig. 1A**). Ca positive samples also had higher quantities of GBS than Ca negative samples (**Fig. 1B**). Together, these data demonstrate that colonization by Ca is correlated with an increase in both the incidence of GBS carriage as well as GBS burdens in pregnant individuals.

**Figure 1:**
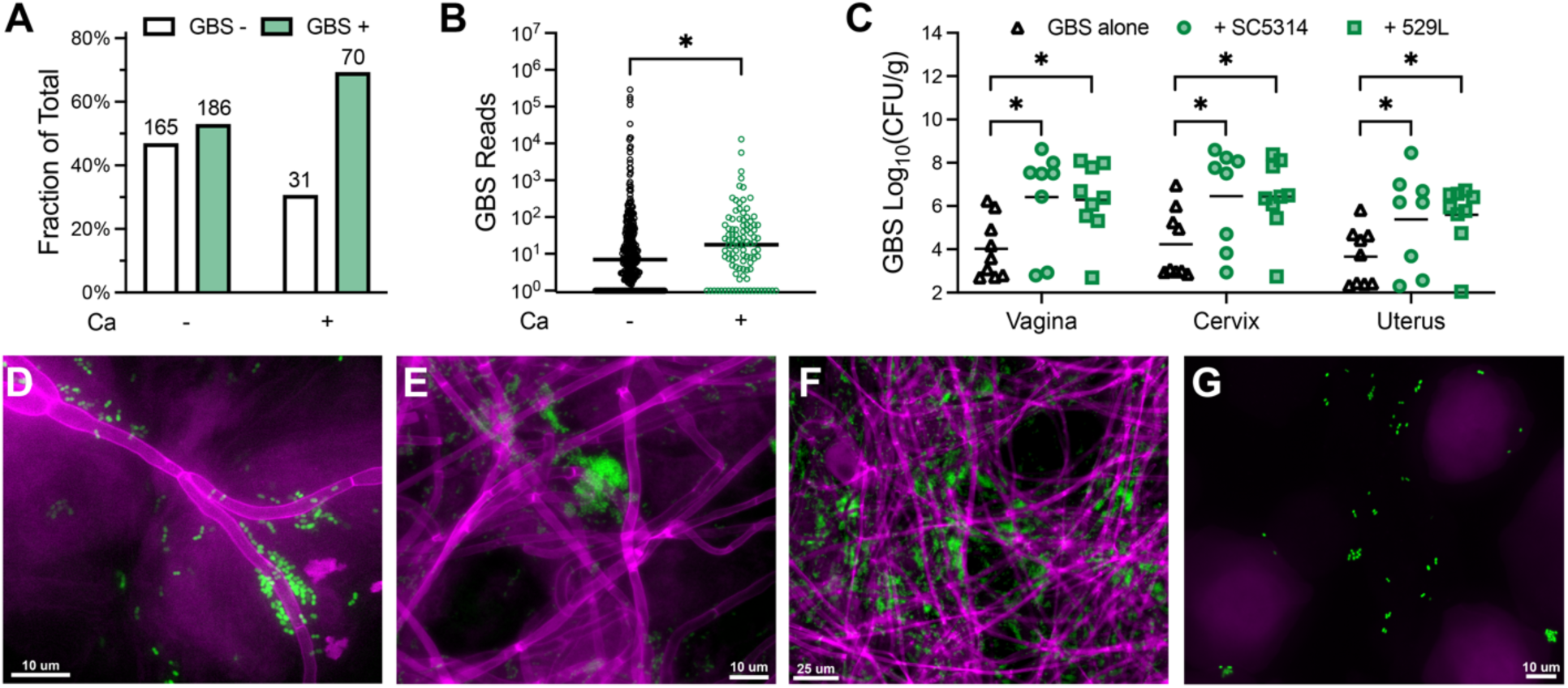
Ca promotes GBS FGT persistence. (A-B) Human vaginal metagenomics, raw data from Baud 2023. **(A)** Proportions of Ca and GBS negative and positive samples. Positive samples had ≥ 5 reads aligned to the indicated organism. Number of samples in group indicated. p = 0.0044, OR = 2.003 (95% CI 1.253-3.189). **(B)** GBS reads in the Ca negative and positive populations. Pseudocount of 1 added to allow for log scale. Medians shown. n = 101-351 samples per group. **(C)** GBS (COH1) burdens four days following GBS intravaginal inoculation. Mice were co-colonized with a mock treatment or Ca (SC5314 or 529L). Means shown. n = 8-9 mice per group. **(D-F)** Vaginal lavage from mice co-colonized with Ca (SC5314; magenta) and GBS (GFP-COH1; green) collected one (D) or two (E-F) days after GBS inoculation. **(G)** Vaginal lavage from mice colonized with GBS alone (GFP-COH1; green) collected two days after GBS inoculation. p values determined by Fisher’s exact test with Baptista-Pike method (A), Mann-Whitney test (B), and one-way ANOVA with Holm-Sidak’s multiple comparisons test (C). * p<0.05

We developed a murine model of vaginal co-colonization to interrogate the effects of Ca on GBS in this host niche as described in the Materials and Methods section. Female CD1 mice were intravaginally inoculated with GBS and either Ca or a mock treatment, and at the experimental end point genital tract tissues were homogenized to quantify microbial loads. To explore whether diverse Ca isolates exhibited interactions with GBS, two genetically and phenotypically divergent strains were utilized: SC5314 and 529L, belonging to clades 1 and 16 respectively^34,35^.

Colonization by either of these two strains significantly increases GBS burdens in the vagina, cervix, and uterus (**Fig. 1C**). While approximately 50% of the mono-colonized animals clear GBS from the genital tract, nearly all co-colonized mice retain GBS in the three tissues. Both Ca strains enhance GBS persistence and uterine ascension to similar degrees. Previous studies have identified that 529L is better able to colonize the murine genital and gastrointestinal tracts compared to SC5314^34,36^. In our vaginal co-colonization model, we similarly observe that 529L reaches significantly higher burdens than SC5314 in the vagina, cervix, and uterus (**Fig. S1A**), despite equivalent GBS burdens in these mice.

To visualize Ca and GBS interactions, vaginal lavage was collected from mice co-colonized with Ca and GFP-expressing GBS, then stained with calcofluor white to visually identify fungi. This revealed that GBS forms aggregates on and around Ca hyphae (**Fig. 1D-F**). The GBS clusters are larger when associated with Ca compared to the more dispersed appearance in GBS mono-colonized mice (**Fig. 1G**). We also detect robust filamentation for both strains of Ca in CD1 and C57BL/6 mice colonized with Ca alone, indicating that Ca still forms hyphal networks *in vivo* in the absence of GBS (**Fig. S1B-D**). We did not find that GBS impacted the rate of Ca filamentation (**Fig. S1E**) or that Ca impacted GBS growth rate (**Fig. S1F-G**) in rich cell culture media.

### GBS and Ca form structured polymicrobial communities

Physical association between Ca and other bacteria has been heavily implicated in both synergistic^37^ and antagonistic^38^ interactions. Following the observation of the striking physical association between GBS and Ca *in vivo*, we sought to characterize the nature of these interactions *in vitro*. In co-culture, we observe GBS cells lining the perimeter of Ca yeast and hyphae (**Fig. 2A**). We developed a quantitative co-aggregation assay to enumerate the proportion of the total GBS population that flocculates with Ca (**Fig. S2**). Following Ca filamentation and addition of GBS, the mixture is centrifuged at a speed optimized to sediment only the Ca cells, due to their larger size, and any attached GBS, physically separating the associated and unassociated bacterial populations. The percentage of GBS within the aggregate can be calculated by quantifying the number of bacteria in the unassociated population.

**Figure 2:**
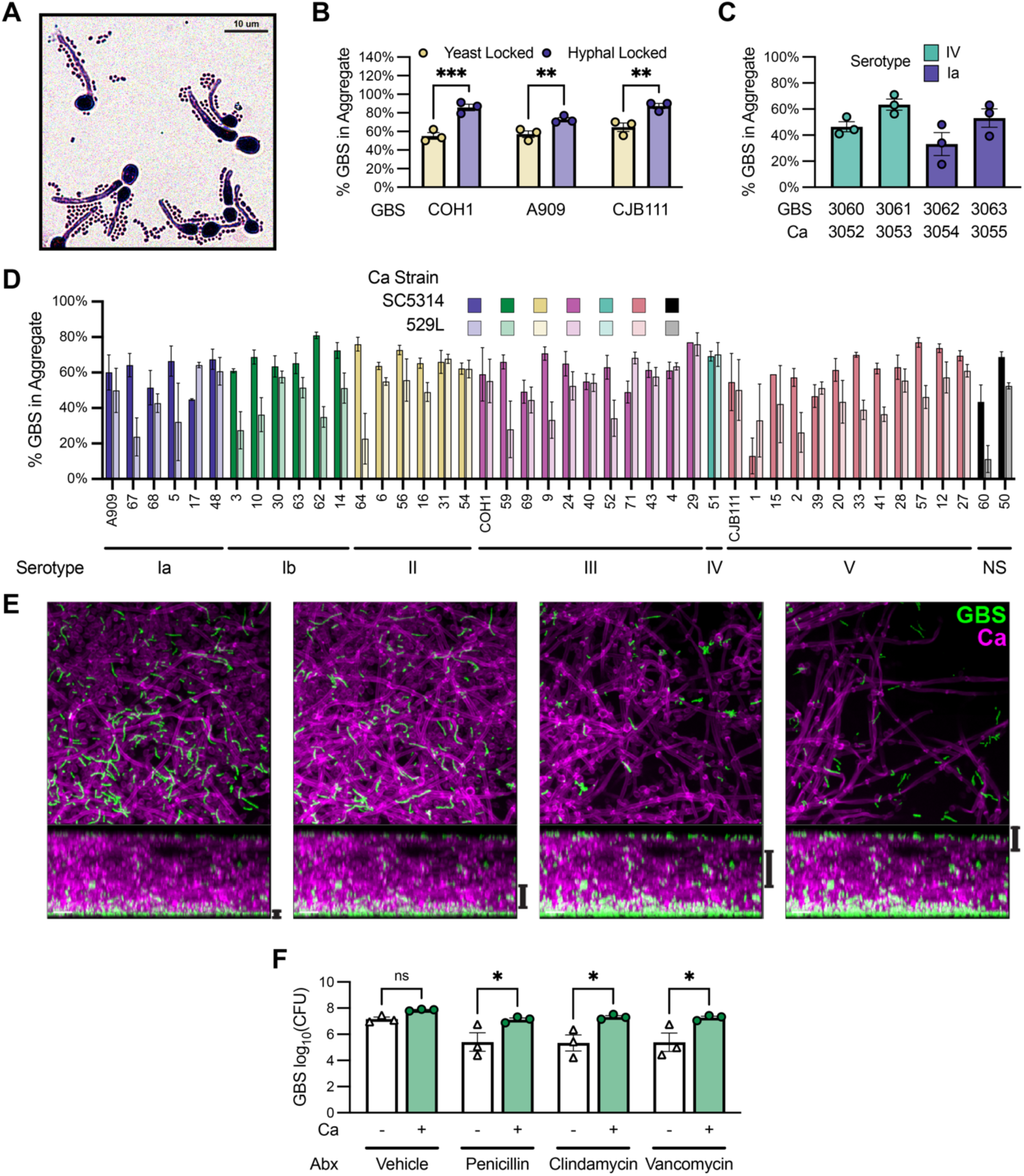
Polymicrobial aggregates protect GBS from antibiotic inhibition. **(A)** Gram stain of GBS (COH1) and Ca (SC5314). **(B)** GBS (COH1, A909, or CJB111) co-aggregation with TetO-*NRG1* Ca. Mean ± SEM from three independent experiments. 12 technical replicates per group. **(C)** Co-aggregation between paired human vaginal co-isolates of GBS and Ca. Mean ± SEM from three independent experiments. 12 technical replicates per group. **(D)** GBS co-aggregation with Ca (SC5314 or 529L). GBS strain identifier and capsular serotype are indicated. NS = not serotyped. Mean ± SD. 4-12 technical replicates per group. **(E)** GBS (GFP-COH1; green) and Ca (SC5314; magenta) in a 48-hour biofilm. Top row shows maximum intensity projections of the z positions indicated by the black bars along the bottom row side-view. **(F)** GBS (COH1) and Ca (SC5314) 24-hour biofilms were treated with a vehicle control, 2 μg/mL penicillin, 2 μg/mL clindamycin, or 4 μg/mL vancomycin for 24 hours, washed, dissociated, and plated for GBS CFU enumeration. Mean ± SEM from three independent experiments. 9 technical replicates per group. p values determined by multiple t tests with Holm-Sidak’s multiple comparisons test. * p<0.05; ** p≤0.01; *** p≤0.001

To assess morphotype-specific differences in GBS binding, we utilized a strain of Ca encoding the hyphal repressor *NRG1* under the control of the repressible TetO promoter^39^. In the presence of anhydrotetracycline (aTC), *NRG1* expression is suppressed, locking Ca in the hyphal morphotype. In the absence of aTC, *NRG1* overexpression inhibits filamentation. While three GBS clinical disease isolates all aggregate at high levels with yeast-locked Ca, aggregation is further increased with hyphal-locked Ca (**Fig. 2B**). We were then interested in whether strains adapted to the vaginal environment would interact similarly. To assess this, we utilized four GBS-Ca strain pairs that were co-isolated from vaginal swabs taken from healthy, non-pregnant individuals. All four pairs form robust aggregates (**Fig. 2C**), supporting that Ca and GBS strains co-isolated from the same vaginal sample are capable of physically co-aggregating. We then screened a second collection of GBS vaginal clinical isolates^40^, along with the three GBS disease isolates COH1, A909, and CJB111, for rates of co-aggregation (**Fig. 2D**). These isolates represent six characterized capsular serotypes and ten sequence types of GBS. 92% of the strains tested exhibit high rates of co-aggregation, above 50%, for either SC5314 or 529L, with a mean of 55% across all strain combinations. 84% of the GBS isolates aggregate at higher levels with SC5314 compared to with 529L, possibly due to the higher rates of filamentation *in vitro* (**Fig. S1E**). Thus, hyphae are potent drivers of co-aggregation, which is strongly conserved across GBS vaginal isolates.

Hyphae are well known to be essential for biofilm formation^41,42^, so we examined whether GBS would colocalize with Ca in this highly organized multicellular community. In the most basal region of the polymicrobial biofilm, the budding yeast morphotype of Ca dominates and GBS is densely clustered with significant chaining between bacterial cells. Moving towards the apical portion of the biofilm, we observe more filamentous Ca hyphae and GBS clusters that wrap around the hyphae. In the most apical region, GBS forms a sparse layer on top of the Ca (**Fig. 2E, Video S1**). An important hallmark of biofilm growth is antimicrobial resistance^43–45^, leading us to test the cultures for inhibition by three clinically relevant antibiotics. Additionally, Ca has been shown to play a role in reducing the efficacy of some antibiotics against GBS strain 515 in planktonic culture and in a zebrafish model of systemic co-infection^46^. Penicillin, clindamycin, and vancomycin represent first, second, and third-line drugs, respectively, that are used during intrapartum antibiotic prophylaxis to prevent GBS transmission to a neonate^47^.

Ca reduces antibiotic inhibition of GBS in all three drug treatments (**Fig. 2F**). This suggests that GBS and Ca can form polymicrobial biofilms together that can protect GBS from antibiotic killing.

### Contact with Ca hyphae increases GBS association to the epithelium

Adherence to the vaginal epithelium is critical for GBS colonization and pathogenesis. To model this host environment, we used hVECs cultured at an air-liquid interface (ALI). This method stimulates the development of a highly differentiated multilayer representative of the vaginal stratified epithelium^48^. Microscopy reveals the layers of epithelial cells as well as the localization of Ca and GBS on and within the cell layers (**Fig. 3A-E; Video S2**). Cultures were inoculated with Ca first to allow for filamentation to occur, followed by inoculation with GBS. Bacterial association to hVEC ALI cultures was assessed by performing a short incubation and extensively washing to remove any non-adherent microbes. GBS is able to adhere to the hVEC ALI cultures on its own, and this adherence is dramatically improved by the presence of Ca, quantified both by microscopy (**Fig. 3F**) and by CFU enumeration (**Fig. 3G**). Ca strains SC5314 and 529L are equally capable of improving GBS association to hVECs. Filamentation is required for this improvement, as yeast-locked Ca does not alter GBS association to the host cells (**Fig. 3H**).

**Figure 3:**
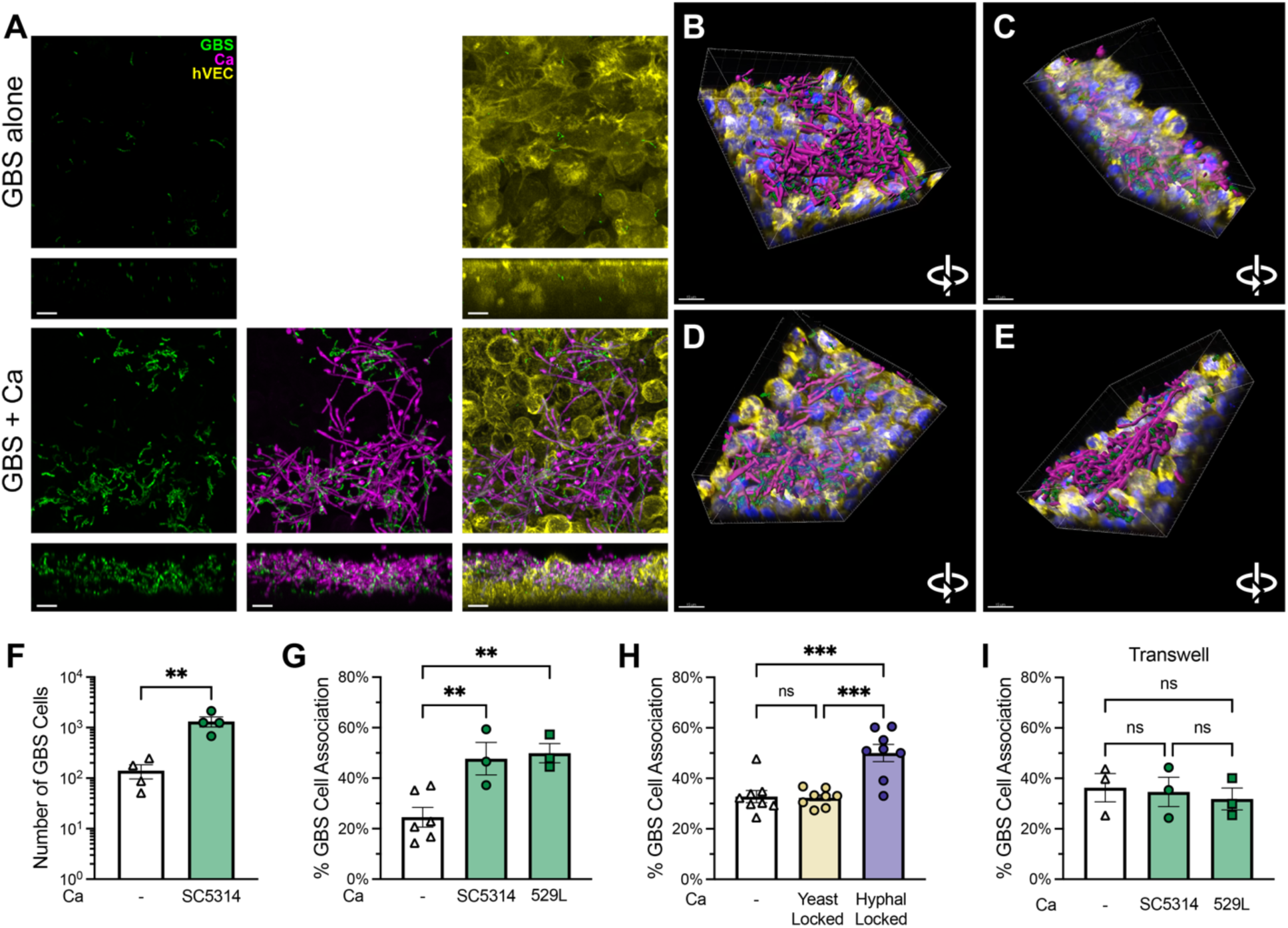
Physical interactions with Ca anchor GBS to host epithelial cells. (A-F) hVEC ALI culture inoculated with GBS (GFP-COH1; green) alone or with Ca (iRFP-SC5314; magenta). **(A)** Cultures stained with Phalloidin-iFluor555 (yellow) to visualize F-actin. Maximum intensity projections and side views are shown. **(B-E)** Stills from Video S2. Volume rendering of GBS (green) and Ca (magenta) with F-actin (yellow) and hVEC nuclei (DAPI; blue). **(F)** Adherent GBS (COH1) on hVEC ALI cultures quantified by confocal microscopy. Mean ± SEM from two independent experiments. 10-12 images per condition. **(G-H)** Adherent GBS (COH1) on hVEC ALI cultures quantified by CFU enumeration. Mean ± SEM from six (G) or eight (H) independent experiments. 9-24 technical replicates per group. **(I)** Adherent GBS (COH1) on hVEC monolayers quantified by CFU enumeration. A transwell insert was placed in each well with either a mock treatment or Ca added to the apical compartment. Mean ± SEM from three independent experiments. p values determined by t test (F) or one-way ANOVA with Holm-Sidak’s multiple comparisons test. * p<0.05; **p≤0.01; ***p≤0.001

Ca adherence to hVEC ALI cultures, calculated by the quantification of Ca DNA present on washed cultures, was not impacted by the presence of GBS (**Fig. S3A**). To investigate whether the extended incubation with Ca alters how permissive hVECs are to GBS adherence, filamentous Ca was added to hVEC ALI cultures at the same time as GBS. Bacterial adherence is significantly improved by the simultaneous addition of Ca and GBS (**Fig. S3B**), indicating that prior and concurrent Ca inoculation are both able to increase GBS attachment to the epithelium. This increase is dependent on contact between Ca and GBS, as separating the microbes by a transwell insert (**Fig. 3I**) and replacing live Ca cells with conditioned medium (**Fig. S3C**) eliminates the Ca-dependent enhancement of GBS adherence. Together, these data suggest that the presence of Ca hyphae can considerably increase the ability of GBS to attach to the vaginal epithelium.

### GBS induces fungal arginine biosynthesis

With evidence suggesting that GBS and Ca co-colonization frequently occurs and can support GBS survival and pathogenesis both in the vaginal lumen and at the epithelial surface, we sought to dissect the host and microbial responses driving these outcomes. To accomplish this, we performed transcriptomic analysis. GBS, Ca, and hVEC ALI monocultures and co-cultures were incubated for six hours, after which RNA from all three organisms was isolated and sequenced. Reads from each mono-and co-culture were aligned to the genomes of the organisms contained in that culture. For co-cultures, only the unmapped reads from each sequential alignment were used as input for alignment to the genome of the next most abundant organism (**Fig. 4A**).

**Figure 4:**
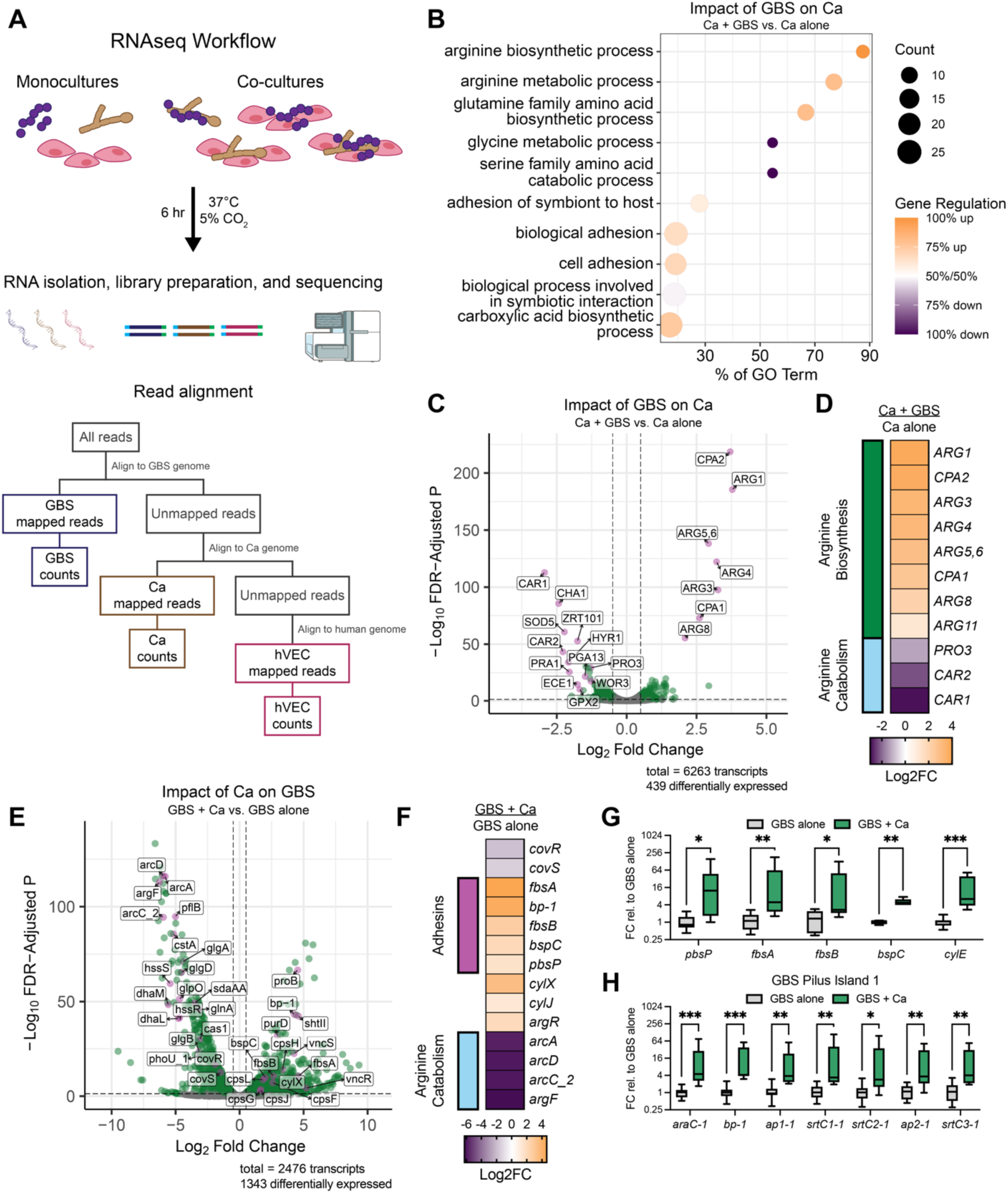
GBS and Ca influence transcription of metabolic and virulence factors. **(A)** Experimental and bioinformatic workflow. GBS (COH1), Ca (529L), and hVEC ALI cultures were co-incubated for six hours. RNA was isolated and sequenced, then reads were mapped to the appropriate genomes. **(B)** Ca differentially expressed (DE) genes with GBS compared to Ca monoculture were analyzed for enrichment of gene ontology (GO) terms. Dot size represents the number of DE genes within a GO term. X-axis represents the proportion of all genes within a GO term that are DE. Dot color represents the proportion of DE genes within a GO term that are up or downregulated. **(C-D)** Ca gene expression with GBS compared to Ca monoculture. **(E-F)** GBS gene expression with Ca compared to GBS monoculture. **(G-H)** GBS (COH1) was grown with Ca (SC5314) for six hours. RNA was isolated and used for RT-qPCR to assess the quantity of the indicated transcripts. 16S rDNA was used as a housekeeping gene. ΔΔCt compared to GBS monoculture. Min to max boxplots shown. Data represent three independent experiments. 5-8 technical replicates per group. p values determined by multiple Mann-Whitney tests. * p<0.05; ** p≤0.01; *** p≤0.001

Gene Set Enrichment Analysis (GSEA) reveals that, in response to GBS, amino acid metabolism is enriched in the Ca differentially regulated gene set **(Fig. 4B**).

Looking at specific genes that are altered, we find that the Ca transcriptional response to GBS is very specific and restricted, with a notable upregulation of nearly every component of the arginine biosynthetic pathway (**Fig. 4C-D**). Exposure to GBS causes the upregulation of Ca arginine biosynthesis and downregulation of the arginine catabolic genes *CAR1*, *CAR2*, and *PRO3*, encoding arginase, ornithine aminotransferase, and pyrroline-5-carboxylate reductase. This strongly suggests that GBS drives arginine biosynthesis and inhibits its breakdown within the Ca cells.

We also find that GBS modulates the expression of several notable fungal virulence factors. This is evidenced by the downregulation of *PRA1*, *ZRT101*, and *ECE1*, encoding a proinflammatory zinc scavenger, its cognate zinc transporter, and the polyprotein from which the cytolytic toxin, candidalysin, is derived (**Fig. 4C**). This suggests that in response to GBS, Ca may be dampening its pathogenic potential. This is in contrast to the upregulation of several GBS virulence factors induced by the presence of Ca. We observe that exposure to Ca drives the upregulation of GBS adhesin genes including *pbsP*, *fbsA*, *fbsB*, *bspC*, and *bp-1* (**Fig. 4E-F**). These encode a plasminogen-binding protein (PbsP), two fibrinogen-binding proteins (FbsA and FbsB), the antigen I/II family Group B streptococcal surface protein C (BspC), and the backbone protein of pilus island 1. Further, Ca induces the downregulation of the two-component system encoded by *covRS*, which represses the expression of a large number of GBS virulence factors and is heavily implicated in streptococcal pathogenesis. Some of the genes repressed by CovRS include the *cyl* operon, which encodes the GBS hemolysin/cytolysin (β-H/C, hemolytic pigment), as well as *pbsP* and *bp-1*. These genes are upregulated in the presence of Ca, which could be associated with the decrease in *covRS* expression. Among the most significantly downregulated genes are *arcD*, encoding the arginine/ornithine antiporter, and *arcA*, *arcC*, and *argF*, which are involved in arginine catabolism. The repressor of this system, encoded by *argR*, is upregulated. Downregulation of arginine import and catabolism suggests that GBS is exposed to higher levels of arginine in co-culture with Ca than in monoculture.

To confirm some of the RNA sequencing results and assess whether these transcriptional changes are strain-specific, we utilized RT-qPCR to quantify GBS gene expression in co-culture with Ca. These results indicate that Ca strain SC5314 is able to induce GBS expression of *pbsP*, *fbsA*, *fbsB*, *bspC*, *cylE*, and the genes encoded on pilus island 1 (**Fig. 4G-H**). Together, transcriptomic evidence suggests that GBS boosts Ca arginine biosynthesis and that Ca may be concurrently increasing GBS pathogenic potential.

### GBS exploits hyphal adhesion at the epithelial surface

Ca and GBS interact with each other in the vaginal lumen, but both organisms are also capable of penetrating the mucus layer and directly contacting the vaginal epithelial cells. To gain a better understanding of whether the epithelium influences these interactions, we investigated how the addition of hVECs alters Ca transcription in culture with GBS. We find that hVECs drive the upregulation of carbohydrate uptake genes *HGT10* and *HGT12* and the downregulation of putative ammonium and ammonia transporters *ATO1*, *FRP6*, and *FRP3* (**Fig. 5A**). Further, Ca upregulates a number of surface-anchored factors that can enable colonization and adhesion to host-derived substrates. Many of these adhesins are strongly induced by the presence of hVECs, and their expression is minimally altered by further addition of GBS (**Fig. 5B)**. These include *HYR1* and *ALS1*, which are among the repertoire of adhesins positively regulated by the transcription factor Bcr1. We find that *BCR1* is required for Ca to promote GBS adherence to hVEC ALI cultures, as a deletion mutant does not increase the number of associated bacteria (**Fig. 5C**). Both *HYR1* and *ALS1*, which are upregulated in proximity to hVECs, contribute to GBS adherence (**Fig. 5D**). *ALS3*, which is not differentially regulated by the presence of hVECs but is induced by Bcr1, also promotes GBS adherence, which has been observed in previous studies^25^ (**Fig. 5D**).

**Figure 5:**
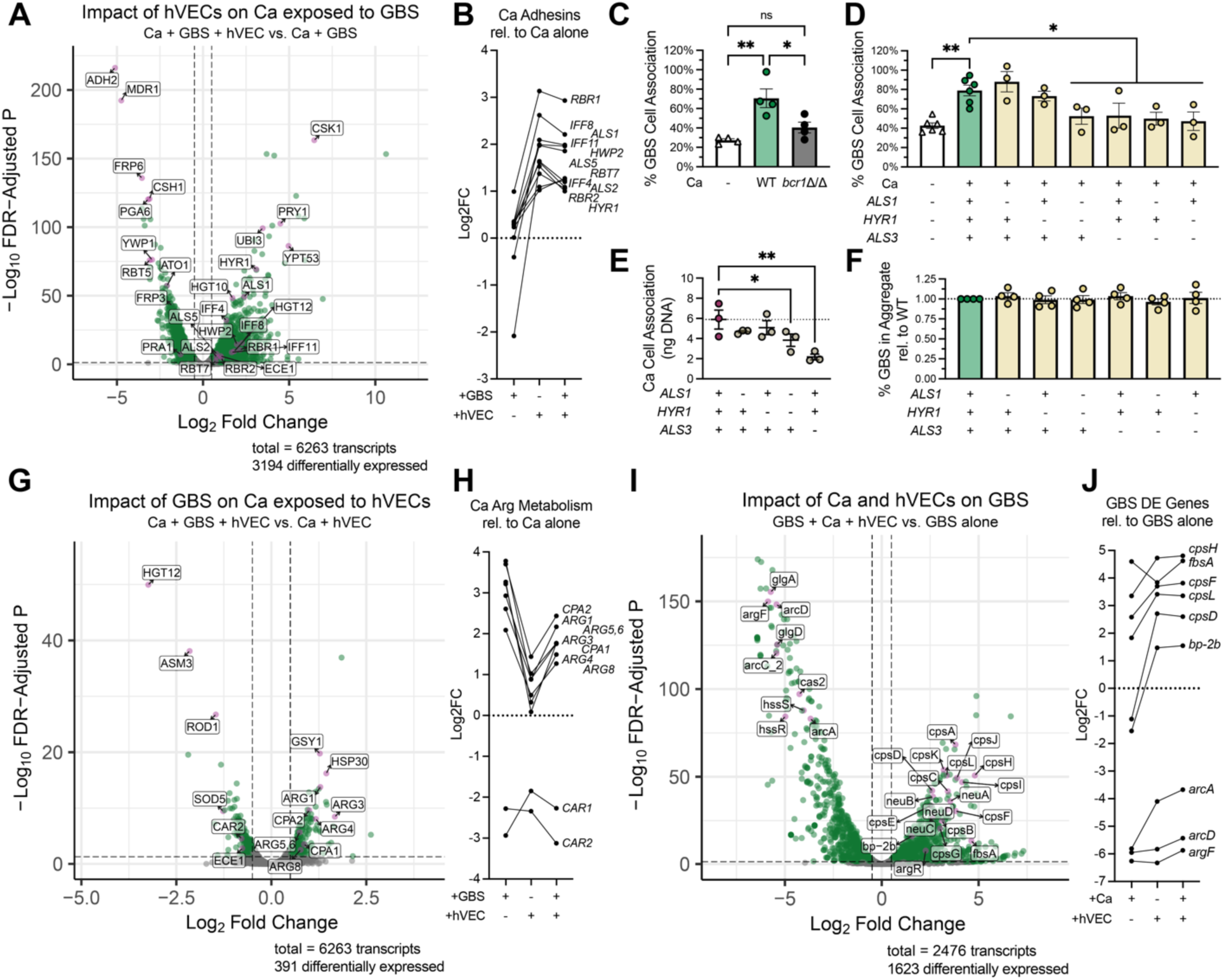
Triple RNAseq reveals transcriptional perturbations during co-culture. **(A)** Ca DE genes with GBS and hVEC ALI culture compared to Ca-GBS co-culture. **(B)** Log2(Fold Change) of select Ca genes in Ca-GBS co-culture, with hVEC ALI culture, or with GBS and hVEC ALI culture compared to Ca monoculture. **(C-D)** Adherent GBS (COH1) on hVEC ALI cultures alone or with Ca (WT or homozygous single or double mutant as indicated) quantified by CFU enumeration. Mean ± SEM from four (C) or six (D) independent experiments. 9-18 technical replicates per group. **(E)** Adherent Ca (WT or homozygous single or double mutant as indicated) quantified by qPCR for Ca 18S rDNA. Mean ± SEM from three independent experiments. 9 technical replicates per group. **(F)** GBS (COH1) co-aggregation with Ca (WT or homozygous single or double mutant as indicated). Mean ± SEM from four independent experiments. 16 technical replicates per group. **(G)** Ca DE genes with GBS and hVEC ALI culture compared to Ca with hVEC ALI culture. **(H)** Log2(Fold Change) of select Ca genes in Ca-GBS co-culture, with hVEC ALI culture, or with GBS and hVEC ALI culture compared to Ca monoculture. **(I)** GBS DE genes with Ca and hVEC ALI culture compared to GBS monoculture. **(J)** Log2(Fold Change) of select GBS genes in Ca-GBS co-culture, with hVEC ALI culture, or with Ca and hVEC ALI culture compared to GBS monoculture. p values determined by one-way ANOVA. * p<0.05; ** p≤0.01

Deletion of both *HYR1* and *ALS1* or deletion of *ALS3* is sufficient to reduce Ca adherence to hVEC ALI cultures (**Fig. 5E**) but does not impact co-aggregation with GBS (**Fig. 5F**). We find that Hyr1 and Als1 contribute to the Ca-dependent enhancement of GBS adherence to hVECs, but that they are not required for co-aggregation with Ca, indicating that the increased adherence may not be attributed to reduced physical interactions between the two microbes. This demonstrates that GBS takes advantage of the presence of Ca and its specialized surface proteins that are induced by the presence of hVECs and enable attachment to host epithelia.

We then addressed the influence of GBS on Ca in the presence of hVECs.

Interestingly, we find that GBS drives down the expression of *HGT12*, in contrast to the hVECs driving up its expression (**Fig. 5G**). Further, GBS stimulates the expression of genes in the arginine biosynthetic pathway similarly to what we observed in the absence of hVECs (**Fig. 4C-D**). We find that arginine biosynthesis is specifically induced by GBS independently of their proximity to hVECs (**Fig. 5H**). Looking at the influence of Ca and hVECs on GBS, we observe downregulation of arginine import and catabolism (**Fig. 5I-J**). GBS also strongly upregulates its capsule biosynthesis gene cluster, as well as the expression of several surface proteins that bind to host factors. These include *fbsA* and *bp-2b*, encoding the backbone protein of pilus 2b which binds host mucins^21^ (**Fig. 5I-J**). The expression of capsule biosynthesis machinery and pilus 2b is primarily driven by the presence of hVECs and these genes remain upregulated upon the addition of Ca.

We find that the GBS population associated to the vaginal epithelial cell surface increases Ca arginine biosynthesis and that these GBS cells are upregulating a unique set of surface factors that play a role in host colonization.

### Ca causes a reduction in transcription of epithelial immune factors

Epithelial cells are among the first host cells to respond to mucosal infection and encode a suite of immune modulatory pathways. In line with this, the hVEC ALI cultures have a strong transcriptional response to incubation with GBS. Some of the most highly upregulated genes include *IL1B*, *TNF*, *NFKB1*, *NFKB2*, *CXCL1*, *CXCL2*, *CXCL3*, and *CXCL8*, all involved in initiating and propagating NF-ΚB signaling (**Fig. S4A**). The hVEC response to Ca is markedly different, with a notable absence of strong pro-inflammatory signaling. Rather, Ca induces barrier signaling and maintenance (**Fig. S4B**). This is evidenced by the upregulation of genes including *OCLN* and *CLDN1*, *2*, and *4*, encoding components of tight junctions, and *EFNA1*, critical for bidirectional epithelial signaling and angiogenesis. We observe wound healing and coagulation associated signaling in response to Ca, demonstrated by the upregulation of *ADM*, *IL24*, *SERPINB2* and *SERPINE1*. A small number of genes involved in antifungal cytokine signaling pathways are also upregulated, albeit at very low levels, including *CLEC7A* and *IL1A*.

We performed GSEA on the genes that hVEC ALI cultures differentially express when exposed to GBS and find that genes involved in the NF-ΚB pathway, immune cell differentiation and activation, and cytokine signaling are enriched in this dataset (**Fig. S4C**). These genes are primarily upregulated upon GBS exposure, further indicating that vaginal epithelial cells initiate pro-inflammatory signaling pathways. We were then interested in investigating the impact of Ca on GBS-exposed hVECs. GSEA on this differentially expressed gene set reveals that the same inflammation-associated ontology terms are enriched in this dataset; however, the genes are nearly all downregulated by the addition of Ca to hVECs with GBS (**Fig. S4C**). Included in this downregulated gene set are the neutrophil chemoattractant *CXCL1*, pro-inflammatory signaling molecule *TNF*, and granulocyte and monocyte activator *CSF2* (**Fig. S4D**).

These genes are not induced by Ca but are upregulated upon GBS exposure. The response is dampened when both Ca and GBS are added to the hVEC ALI cultures (**Fig. S4E-F**). Together, this suggests that Ca downregulates the host antimicrobial response to GBS, indicating a possible role for fungal colonization in GBS immune evasion.

### Arginine induces GBS factors that contribute to synergy with Ca

Transcriptomics analysis revealed that GBS induces Ca arginine biosynthesis, with *ARG1* being the most highly upregulated gene in the biosynthetic pathway upon GBS exposure. To further interrogate this, we utilized a reporter strain of Ca expressing GFP under the *ARG1* promoter. GFP expression increases significantly in co-culture with GBS (**Fig. 6A-B**). Further, all four GBS vaginal isolates that were co-isolated with Ca strongly induce *ARG1* promoter activity. We find that this is a contact-dependent phenotype, as separating GBS from Ca with a transwell insert eliminates the increase in *ARG1* promoter activity. Referring to the GBS clinical isolate co-aggregation screen (**Fig. 2D**), we compared the inducing effects of the isolate with the lowest level of co-aggregation, isolate 1, to the highest, isolate 62. Isolate 62 induces *ARG1* at significantly higher levels than isolate 1 (**Fig. 6B**). Taken together, these results suggest that GBS-Ca physical interaction is required for the induction of Ca arginine biosynthesis.

**Figure 6:**
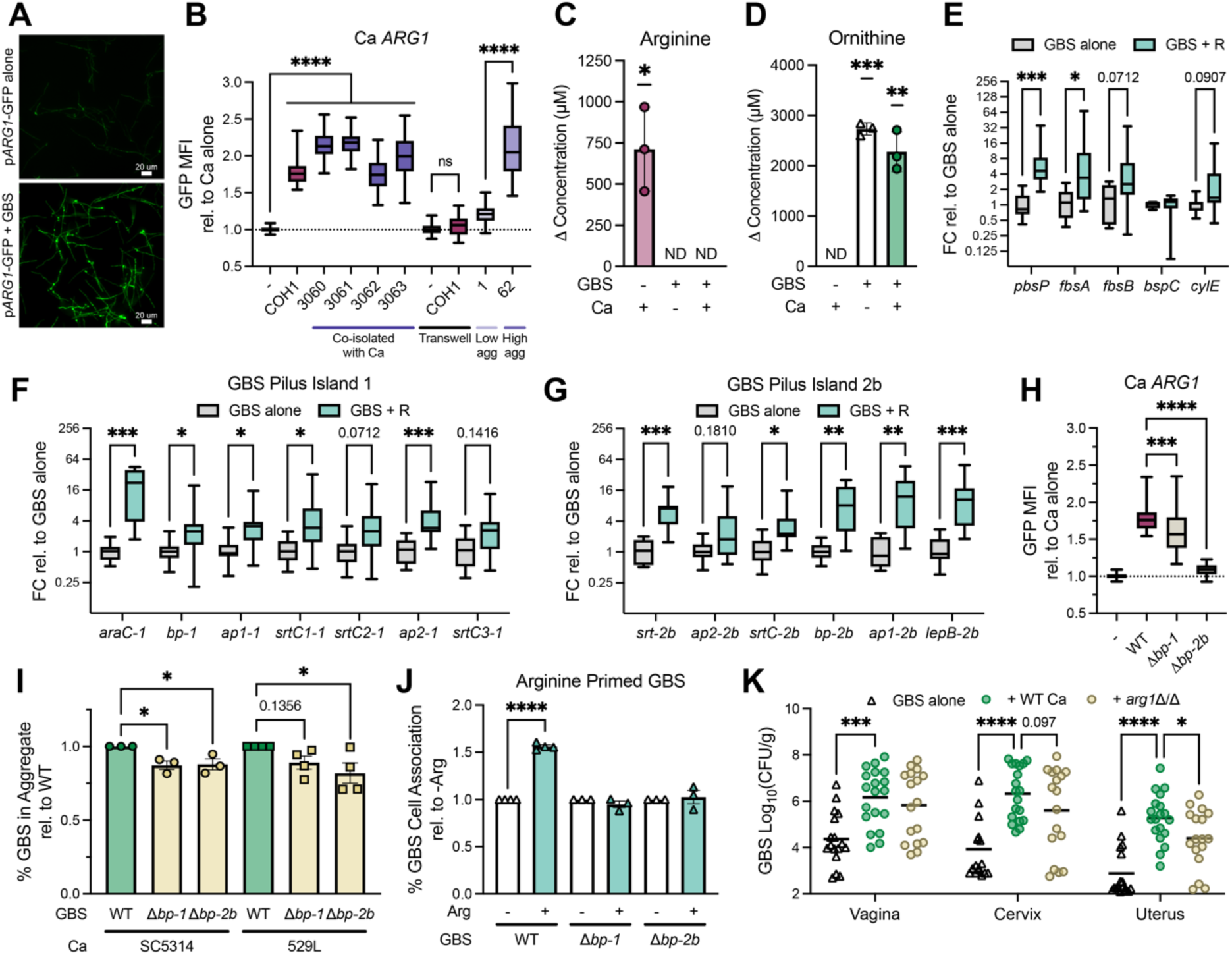
Ca arginine biosynthesis increases GBS pathogenic potential. **(A)** Ca *ARG1* reporter (p*ARG1*-GFP p*ADH1*-yCherry; green) alone or GBS (COH1). GFP channel is shown. **(B)** Ca *ARG1* reporter incubated alone or with the indicated GBS strain. GBS was separated from Ca by a transwell insert where indicated. GFP mean fluorescence intensity (MFI) was calculated within the area of yCherry fluorescence. GFP MFI relative to Ca monoculture is shown. Min to max boxplots from three independent experiments. 36-72 images per condition. **(C-D)** Ca (SC5314), GBS (COH1), or both were grown for six hours. Arginine (C) and ornithine (D) concentrations quantified by UPLC. Baseline media concentration, or LOD for values too low to detect, subtracted. Stars represent comparisons to zero. Arginine in GBS monoculture and GBS-Ca co-culture, and ornithine in baseline media and Ca monoculture were below the LOD of 3 μM. ND = not detected. Mean ± SD. 3 technical replicates per group. **(E-G)** RT-qPCR of GBS (COH1) grown in standard KSFM or KSFM supplemented with 12 mM arginine. 16S rDNA used as a housekeeping gene. ΔΔCt compared to standard media. Min to max boxplots from three independent experiments. 5-9 technical replicates per group. **(H)** Ca *ARG1* reporter incubated alone or with GBS (COH1 WT or mutant). GFP MFI was calculated within the area of yCherry fluorescence. GFP MFI relative to Ca monoculture is shown. Min to max boxplots from three independent experiments. 36-72 images per condition. **(I)** GBS (COH1 WT or mutant) co-aggregation with Ca (SC5314 or 529L). Mean ± SEM from four independent experiments. 12 technical replicates per group. **(J)** GBS (COH1 WT or mutant) overnight cultures grown in KSFM were diluted into standard KSFM or KSFM supplemented with 12 mM arginine. At mid-log, GBS was inoculated onto hVEC ALI cultures and incubated for 30 minutes. Cultures were washed, dissociated, and plated for CFU enumeration. Mean ± SEM from four independent experiments. 11-14 technical replicates per group. **(K)** GBS (COH1) burdens four days after GBS intravaginal inoculation. Mice were co-colonized with a mock treatment or Ca (SC5314 WT or *arg1*Δ/Δ). N = 17-19 mice per condition. p values determined by one-way ANOVA (B, H-J), one sample t test (C-D), multiple Mann-Whitney tests (E-G), or two-way ANOVA (K) with Holm-Sidak’s multiple comparisons test. * p<0.05; ** p≤0.01; *** p≤0.001; **** p≤0.0001

GBS does not encode the complete arginine biosynthetic pathway, making it an arginine auxotroph^49^. To investigate whether Ca could supply this essential metabolite to GBS, we measured the amino acid concentrations in media conditioned with Ca and/or GBS. The Ca monoculture significantly increases arginine concentrations above baseline, indicating that it is capable of secreting it into the media (**Fig. 6C**). GBS depletes the arginine levels to below the limit of detection, highlighting that it can import and potentially utilize millimolar levels of arginine within a six-hour time period. GBS imports arginine using the ArcD arginine-ornithine antiporter, and we observe that GBS significantly increases the levels of ornithine in the media, concordant with high levels of antiporter activity (**Fig. 6D**). Without arginine, GBS does not exhibit any measurable growth. Supplementation with 10% WT Ca conditioned medium is sufficient to restore growth to the extent seen in arginine replete conditions; however, conditioned medium from *arg1*Δ/Δ or *arg3*Δ/Δ Ca does not restore growth to the same level (**Fig. S5A**).

Because arginine exposure has been associated with increasing GBS hemolytic activity^50^, we hypothesized that it could impact other facets of its pathogenic potential. This led us to ask whether arginine nutrient sharing may be involved in priming GBS by upregulating virulence factors that support colonization. We find that the addition of supplemental arginine is sufficient to increase transcription of *pbsP* and *fbsA* (**Fig. 6E**), in addition to pilus islands 1 and 2b (**Fig. 6F-G**). Interestingly, both pili are involved in stimulating *ARG1* expression (**Fig. 6H**) and contribute to co-aggregation with Ca (**Fig. 6I**). The arginine biosynthetic pathway; however, is not required for GBS and Ca to co-aggregate, indicating that other factors are also involved and that more incubation time may be necessary for Ca to synthesize and secrete arginine (**Fig. S5B**). We find that growing GBS in media containing supplemental arginine boosts adherence to hVEC ALI cultures by 50%, and this priming effect is dependent on the presence of both pili (**Fig. 6J**). This supports a model in which arginine-induced upregulation of pili 1 and 2b increases GBS adhesion to host surfaces. These data outline a positive feedback loop in which the same factors that contribute to physical interactions between GBS and Ca also drive arginine biosynthesis, which in turn upregulates bacterial surface proteins that increase adhesion and virulence potential.

Having shown that excess arginine can prime GBS for increased adhesion to the epithelium, we hypothesized that Ca arginine biosynthesis plays a role in bacterial persistence during co-colonization. To test this, mice were colonized with GBS and either WT Ca, *arg1*Δ/Δ Ca, or a mock treatment. Genital tract tissues were collected four days following GBS inoculation. Co-colonization with WT Ca increases GBS burdens in the vagina, cervix, and uterus. Deletion of *ARG1* causes a defect in the ability of Ca to support GBS, trending in the cervix and statistically significant in the uterus (**Fig. 6K**).

This is not due to a difference in Ca burdens as WT and *arg1*Δ/Δ colonize the vagina at equivalent levels throughout the duration of the experiment (**Fig. S5C**). GBS co-colonization with *arg1*Δ/Δ Ca still results in an increase in bacterial burdens compared to mono-colonized mice. This indicates that other fungal factors in addition to arginine biosynthesis contribute to interactions with GBS *in vivo*, with co-aggregation and modulation of the host response possibly among these factors. The Ca arginine biosynthetic pathway is not sufficient to increase GBS persistence in the vagina, but it does play a role in promoting GBS burdens in the upper genital tract tissues.

## Discussion

An emerging body of work describes how the mycobiome significantly impacts human health by modulating the composition and lifestyles of resident microbes at a number of body sites^51^. These interactions have been described primarily for *Candida* species and mostly in the gastrointestinal tract where fungal colonization not only plays a role in maintaining bacterial diversity during homeostasis but is important for overcoming disruptions caused by antibiotics^52^. In this study, we use *in vitro* and *in vivo* models to demonstrate and characterize the strongly conserved physical interactions between Ca and GBS, highlighting how they can lead to multiple beneficial outcomes for GBS. These include evading antibiotic inhibition, adhering to the host epithelium, and accessing arginine, an essential nutrient for GBS which Ca secretes into the extracellular space. Arginine exposure further increases GBS adhesion to the epithelium and stimulates the expression of factors involved in adhesion to Ca itself, resulting in increased arginine biosynthesis by Ca (**Fig. 7**). Ultimately, we find that Ca arginine biosynthesis contributes to GBS ascension to the uterus.

**Figure 7:**
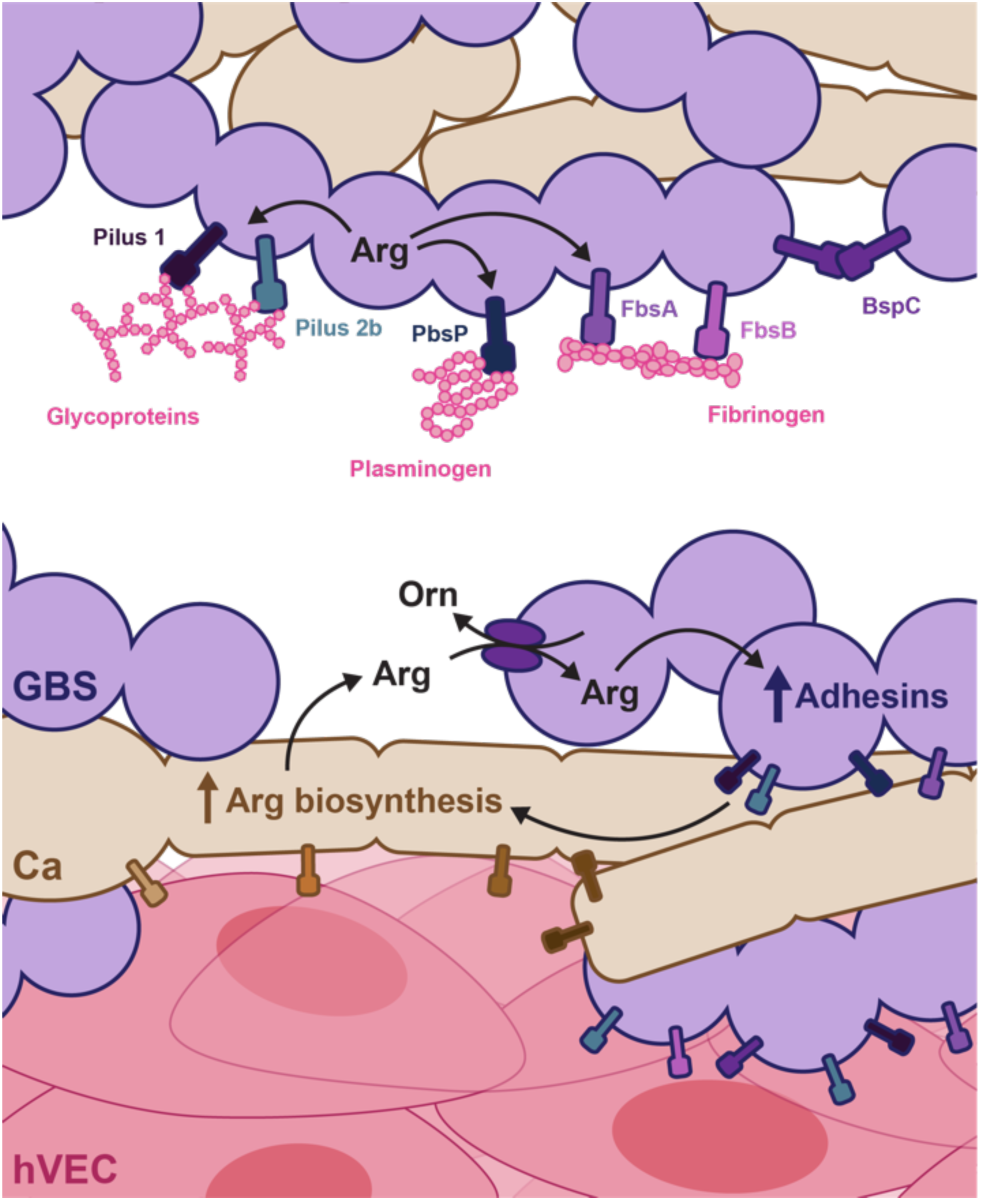
GBS-Ca nutrient sharing leads to a positive feedback loop that increases adhesion. GBS and Ca co-localize in the vaginal lumen and on the surface of vaginal epithelial cells. Physical contact promotes arginine biosynthesis in Ca, which can be secreted and imported into GBS via the arginine-ornithine antiporter. High levels of arginine upregulate adhesin expression. These surface proteins promote GBS adhesion to extracellular matrix components, vaginal epithelial cells, and Ca, further upregulating Ca arginine biosynthesis.

Different *Candida* species have been shown to alter the composition of the vaginal microbiome in unique ways^53^, although further studies are needed to fully characterize how the *Candida*-associated vaginal microbiome might impact GBS vaginal persistence. To interrogate the downstream effects of Ca colonization, we developed a murine model in which mice synchronized in their estrus cycles are intravaginally inoculated with Ca first to establish colonization, then subsequently inoculated with GBS. This model demonstrates that two Ca strains, SC5314 and 529L, can increase GBS burdens across all tissues of the FGT. There are multiple studies highlighting how these two strains differ in the regulation of many cell processes and in their interactions with the host^34,54–56^, demonstrating the importance of strain selection during study design and the wide levels of variance in phenotype within a single species. The conserved phenotype with GBS could indicate that even highly divergent Ca isolates, such as those found in different human populations, might have the capacity to increase GBS survival and persistence in the FGT.

We find that Ca physically interacts with GBS in the vaginal lumen, with clear evidence of their colocalization. GBS self-association is dramatically altered in the lumen of co-colonized mice compared to mice colonized with GBS alone due to the formation of dense aggregates on and around Ca hyphae. Several studies have described bacterial association to Ca and other filamentous fungi including *A. fumigatus*, *A. flavus*, and *R. microsporus*^57^. Some bacteria can migrate along hyphae which promotes dispersal^58,59^ and may facilitate downstream interactions with other microbial species^60^. Physical interactions between Ca and *Streptococcus* species are mediated both by adhesins and by modulation of cell wall components^61^, which also represent critical interfaces for interactions with host substrates.

In this study, hVEC ALI cultures are used to model microbial interactions directly at the vaginal epithelial surface. To identify whether Ca and GBS have different transcriptional responses to each other in the vaginal lumen and at the epithelial surface, we grew the microbes together in cell culture media with or without hVEC ALI cultures. In co-culture, we broadly find that GBS lowers Ca pathogenic potential by downregulating virulence factors *PRA1* and *ECE1*. GBS also induces upregulation of *YWP1* and *ENG1* which control β-glucan masking and shaving respectively, potentially contributing to evasion of recognition by dectin-1^62,63^. Previous work has described GBS decreasing Ca expression of *EFG1* and inhibiting filamentation^64^. While we did not observe either of these in this study, it provides further evidence that Ca virulence may be dampened in the presence of GBS. In contrast, our work suggests that Ca may be increasing GBS pathogenic potential at the transcriptional level. In this study we show that, in co-culture with Ca, many genes in the CovRS regulon are upregulated including pilus island 1 and *pbsP*, which have been shown to bind host glycoproteins^22^ and plasminogen^23^ respectively. The CovRS TCS is a master regulator of virulence, controlling expression of the *cyl* operon which encodes the GBS β-H/C. Ca has been shown to regulate TCSs and global regulators of virulence factors in other organisms including *S. aureus*^65,66^ and *S. mutans*, where the expression of the TCS regulon directly impacts the outcome of these interkingdom interactions^67^.

We demonstrate that hVECs induce unique and robust transcriptional programming in Ca. This includes causing the upregulation of several GPI-anchored surface proteins, many of which are predicted to or have been shown to contribute to adhesion and persistence. Previous studies have shown that the Ca adhesin Als3 is involved in promoting GBS adherence to hVEC and bladder epithelial cell monolayers^25,68^. We similarly observe that Als3 is important for Ca to increase GBS adherence to hVEC ALI cultures, although we do not find that GBS or hVECs induce upregulation of this adhesin. Other surface proteins that are upregulated, Hyr1 and Als1, also contribute to GBS adherence. This could suggest that redundancy across Ca adhesins offers multiple parallel mechanisms to promote GBS adhesion to the host epithelium.

In response to Ca strain 529L, we observe that the host cells initiate wound healing, angiogenesis, and coagulation gene expression signatures. This is in contrast to previous work that has identified pro-inflammatory inflammasome activation as a key host response to other Ca strains, SC5314 and 3153A, which may be representative of symptomatic VVC^69–71^. Coagulation has previously been associated with antifungal defense^72,73^, and immunothrombosis can be initiated as a component of the innate immune response^74,75^ with the risk of transitioning into excessive microthrombosis and embolism.

Among the most striking transcriptional signatures in our dataset, we find that GBS strongly induces arginine biosynthesis in Ca. A small number of studies have collectively begun to describe how bacteria can induce Ca arginine biosynthesis^76,77^ and how arginine mediates an increase in bacterial virulence during co-infection or co-colonization with Ca^78,79^. In the gut, *Salmonella* Typhimurium induces Ca arginine biosynthesis which dampens host antimicrobial responses and induces bacterial invasion and dissemination. Arginine production by Ca within dental plaque is stimulated by *Actinomyces viscosus* and drives the development of polymicrobial biofilms and root caries. Interestingly, studies independent of Ca have shown that exogenous arginine can increase GBS hemolytic activity^50^.

On the hVEC ALI cultures, we still observe GBS driving upregulation of the Ca arginine biosynthetic pathway and downregulation of arginine catabolism. This could indicate that GBS may be utilizing Ca-derived arginine both in the vaginal lumen and at the epithelial surface. Additionally, we observe that Ca induces downregulation of GBS arginine import and the arginine deiminase pathway both in the presence and absence of hVECs. This further supports the hypothesis that GBS is exposed to higher levels of arginine with Ca. Interestingly, previous work has demonstrated that Ca reduces the capability of GBS to acidify culture media^46^. In the presence of excess arginine, GBS can use the arginine deiminase pathway to generate ammonia, causing an increase in pH and enhancing its survival during acid stress. In the case that this pathway is not active, arginine alone will raise the pH as a basic amino acid. Further work is necessary to address whether arginine provided by Ca directly impacts the local pH and through which mechanisms.

Host cells may also be sensing and responding to an increase in local arginine concentrations. Arginine concentrations in the vaginal lumen are estimated to be approximately 200 μm^80^; however, local concentrations are challenging to measure. Concentrations in the Ca-GBS aggregate microenvironment may be very different than bulk measurements are able to quantify. In addition to impacting GBS, arginine can have diverse impacts on the immune system^81^. As a precursor to reactive nitrogen species, arginine can be involved in promoting an inflammatory and highly antimicrobial environment. Conversely, arginase can convert arginine into ornithine which, in macrophages, is indicative of M2 polarization and is associated with tissue repair and proliferation. Arginine availability and metabolism have complex influences on immune cell function and downstream host-microbe interactions^82^. Future studies are required to address if the host is sensing Ca-derived arginine or GBS-derived ornithine, and if these metabolites are altering the environment in a way that could promote GBS vaginal persistence or intrauterine infection.

Our transcriptomics data suggest that Ca may be involved in dampening the antimicrobial immune signaling initiated after GBS inoculation. While further confirmation of these results is required, and is the subject of own ongoing work, previous studies have identified a role for Ca in reducing protective immune responses against other bacteria. Airway colonization by Ca can sensitize the host to bacterial pneumonia^83^, and Ca can suppress pro-inflammatory responses to *F. nucleatum*^84^, *P. gingivalis*^85^, and *S.* Typhimurium^79^. Ca has also been shown to dampen type 17 responses by expression of Lip2^86^, which is upregulated on hVECs in our dataset, both with and without GBS present. Further, type I interferon responses, which are protective against Ca damage to hVECs^87^ and induced by VVC-associated strains^88^, are suppressed by Ca expression of Cmi1, which we find is upregulated both by GBS and by hVECs. Dampening the immune response could not only allow GBS to evade host-mediated killing, but it could also be involved in decreasing the risk of preterm birth despite GBS infiltration into the uterus. A high Th17/Treg ratio has been heavily implicated in triggering preterm labor, in addition to high levels of IL-6, IL-1β, CXCL1, and CSF2^89–91^. Our data suggest a dampening of these type 17 responses during GBS-Ca co-colonization compared to GBS mono-colonization. GBS infection is a risk factor for premature rupture of membranes and preterm labor, and additional work is necessary to assess whether Ca co-colonization may be involved in decreasing these risks.

In this study we describe for the first time a role for Ca-mediated arginine exchange and metabolism in promoting adhesive and pathogenic activity in GBS. These data suggest that targeting interkingdom nutrient sharing could decrease the incidence of GBS ascending infection. Together, we identify arginine as a principal metabolite in the interactions between Ca and GBS, with downstream impacts on the lifestyles and pathogenic potentials of these critical FGT pathobionts.

## Resource availability

### Lead contact

Kelly S. Doran

### Materials availability

Materials will be made available upon request.

### Data and code availability

We do not report any original code in this manuscript. Sequencing data have been submitted to GEO and will be made available upon publication in a peer-reviewed journal.

## Acknowledgements

Quantitative metabolomics was performed by the University of Pennsylvania Microbial Culture and Metabolomics Core. Library preparation and RNAseq was performed at SeqCenter (seqcenter.com, Pittsburgh PA). We thank the Advanced Light Microscopy Core at the University of Colorado Anschutz Medical Campus for their expertise and the University of Colorado Boulder Research Computing Center for assistance with and maintenance of computing services. We thank our collaborators for sharing strains and mutants with our group. The Ca adhesin single and double mutant library in the fRS29 background was a kind gift from Dr. Rebecca Shapiro, and the p*ARG1* reporter and arginine biosynthesis mutants were generously provided by Dr. Michael Lorenz. We thank the Doran Lab members for their thoughtful feedback during data collection and manuscript generation. Work is supported by NIH grants R01AI153332 to K.S.D. and R21AI88719 to K.S.D. and K.S.O, and a Human Infection Challenge Network for Vaccine Development (HIC-Vac) grant (MR/R005982/1) funded by the GCRF Networks in Vaccines Research and Development and co-funded by the MRC and BBSRC to K.L.D.

## Author contributions

Conceptualization, S.C, K.S.D; Data curation, S.C; Formal analysis, S.C; Funding acquisition, K.S.O, K.S.D, K.L.D; Investigation, S.C, A.J.C; Methodology, S.C; Project administration, K.S.D; Resources, M.N.N, K.L.D, A.H.N, K.S.O, K.S.D; Supervision, K.S.O, K.S.D; Visualization, S.C; Writing – original draft, S.C; Writing – review and editing, all authors.

## Methods

### Paired vaginal isolate collection

Clinical co-isolates were collected as part of the TIMING (opTImisation of Methods for a human INfection model for Group B *Streptococcus*) study led by K.L.D (https://cnpi-vaccinology.com/timing/)^92^. Low vaginal Copan Italia SPA swabs were taken from healthy, non-pregnant women aged between 18-40 in the UK and inoculated into Todd-Hewitt Broth with nalidixic acid (15 mg/mL) and colistin (10 mg/mL) (lysine indole motility medium, LIM broth). LIM cultures were incubated for 18 h at 37°C with 5% CO_2_, and then plated onto CHROMagar StrepB or CHROMagar Candida (CHROMagar, Saint-Denis France) to select for GBS or Ca isolates, respectively. The identity of GBS isolates was further confirmed by MALDI-TOF following sub-culture on Columbia horse blood agar. Serotyping of GBS isolates was performed by PCR using extracted genomic DNA as template^92^.

### Strains, cell lines, and media

GBS isolates were grown statically in Todd Hewitt Broth (THB) and plated on Todd Hewitt Agar (THA) at 37°C unless otherwise indicated. *C. albicans* isolates were grown shaking in Yeast Peptone Dextrose (YPD) broth and plated on YPD agar at 30°C unless otherwise indicated. Human vaginal epithelial cell (hVEC) line VK2-E6/E7 was cultured in Keratinocyte Serum-Free Medium (KSFM) supplemented with 0.05 mg/mL bovine pituitary extract (BPE) and 0.5 ng/mL epidermal growth factor (EGF). “KSFM” in this manuscript is used to refer to media containing these concentrations of BPE and EGF. Modified chemically defined media was prepared as described previously^93^ with arginine omitted or added at the indicated concentrations. Biofilms were grown in Brain Heart Infusion (BHI) broth with 10% fetal bovine serum (FBS). For GBS and Ca selection and differentiation, murine lavage and tissue samples were plated on CHROMagar StrepB and CHROMagar Candida. For GBS selection against Ca *in vitro*, samples were plated on THA containing 50 μg/mL nystatin. Conditioned medium for adherence assays (Fig. S3C) was generated by replicating adherence assay conditions: adding 10^6^ cells/mL Ca and/or 10^6^ CFU/mL mid-log GBS in KSFM to the wells of a 96-well plate, incubating for 2.5 hours at 37°C with 5% CO_2_, then filter sterilizing media to be stored at-20°C until use. Conditioned medium for growth curves (Fig. S5A) and metabolite quantification (Fig. 6C-D) was generated by replicating RNA sequencing conditions: adding 2×10^7^ Ca cells/mL and/or 2×10^8^ mid-log GBS CFU/mL in KSFM to the wells of a 6-well plate, incubating for six hours at 37°C with 5% CO_2_, then filter sterilizing media to be stored at-20°C until use.

### Bacterial strains

| Strain | Source |
| --- | --- |
| COH1 | 94 |
| A909 | 95 |
| CJB111 | 96 |
| Paired vaginal isolate collection | 92 |
| Vaginal isolate collection from Detroit Medical Center | 40 |
| COH1 $\Delta bp-1$ | 97 |
| COH1 $\Delta bp-2b$ | 97 |
| COH1 pDESTerm::GFP | 98 |

### Fungal strains

| Strain | Source |
| --- | --- |
| SC5314 | 99 |
| 529L | 36 |
| SN250 | 100 |
| Paired vaginal isolate collection | 92 |
| SC5314 TetO-*NRG1* | 101 |
| SN520 *bcr1*Δ/*bcr1*Δ | 100 |
| fRS29 | 102 |
| fRS29 *als1*Δ/*als1*Δ | 102 |
| fRS29 *hyr1*Δ/*hyr1*Δ | 102 |
| fRS29 *hyr1*Δ/*hyr1*Δ *als1*Δ/*als1*Δ | 102 |
| fRS29 *als3*Δ/*als3*Δ | 102 |
| fRS29 *als3*Δ/*als3*Δ *als1*Δ/*als1*Δ | 102 |
| fRS29 *als3*Δ/*als3*Δ *hyr1*Δ/*hyr1*Δ | 102 |
| SC5314 *arg1*Δ/*arg1*Δ | 103 |
| SC5314 *arg3*Δ/*arg3*Δ | 103 |
| RM1000 pARG1-GFP pADH1-yCherry | 103 |
| SC5314 pENO1-iRFP670 | 101 |

### Metagenomics data analysis

Normalized read counts with assigned taxa were downloaded from Baud et al.^33^ and processed to quantify correlative relationships between the presence of different microbes. Data were filtered to include samples from individuals with at-term births and were binned as positive for a microbe if they contained at least 5 reads assigned to the associated taxon.

### Murine models

8-12 week old female CD1 mice purchased from Charles River Laboratories were used for all experiments unless otherwise indicated. Where indicated, 8-12 week old C56BL/6 mice purchased from Jackson Laboratories were used. CD1 mice were treated with a cocktail of 0.5 mg/mL ampicillin, neomycin, and gentamycin in their drinking water for 7-10 days prior to the experiment. At day-3, mice were synced in their estrus cycles by intraperitoneal injection of 0.5 mg 17β-estradiol dissolved in 100 μL of sterile-filtered sesame oil. The following day (day-2), mice were intravaginally inoculated with 10^7^ yeast cells in 10 μL of PBS. On day-1, mice were given fresh drinking water without antibiotics. On day 0, mice were intravaginally inoculated with 10^7^ mid-log GBS CFU in 10 μL of PBS. The vaginal tract was lavaged using 100 μL of PBS to assess microbial burdens. Lavage was plated on CHROMagar for CFU enumeration. Mice not colonized with GBS one day post GBS inoculation were excluded. At the experimental endpoint, day 4 unless otherwise indicated, animals were sacrificed and female genital tract tissues were harvested, homogenized, and plated on CHROMagar for CFU enumeration. The limit of detection was 40 CFU/mL. If microbial burdens were below the limit of detection, the limit of detection was plotted. Animal experiments were approved by the Institutional Animal Care and Use Committee at the University of Colorado Anschutz Medical Campus on protocol #00316.

### Lavage imaging

Vaginal lavage from mice co-colonized with Ca (SC5314) and GBS (GFP-COH1) was collected by pipetting up and down 3-5 times with 50 μL of PBS twice, generating 100 μL of lavage total. Lavage fluid was centrifuged at 10,000 g for 5 minutes, the supernatant was removed, and the pellet was resuspended in 10 μL of a calcofluor white solution. This suspension was imaged directly on a Keyence BZ-X700 inverted fluorescence microscope equipped with a GFP filter cube (ex 470/40 em 525/50) and a DAPI filter cube (ex 360/40 em 460/50).

### Gram stain and imaging

Ca (SC5314) was incubated in an 8-well glass-bottom chamber slide for 2 hours before adding mid-log GBS (COH1) for an additional 30 minutes in KSFM. The samples were washed with PBS, fixed in 10% paraformaldehyde for 10 minutes, then washed with PBS again. The chambers were removed from the glass slide, leaving any attached microbes. Slides were air-dried then stained with crystal violet and washed, fixed with iodine, washed, decolorized with ethanol and acetone, washed again, and counterstained with fuchsin before a final wash. Slides were imaged on a Keyence BZ-X700 inverted microscope with a color camera.

### Filamentation assay

10^7^ cells/mL of Ca yeast (SC5314 or 529L) and, where indicated, 10^8^ CFU/mL of GBS (COH1) were cultured in KSFM, shaking at 37°C for 1 hour. Samples were mounted on glass slides and ten random locations on each slide were imaged to quantify the number of Ca yeast and the number of cells undergoing filamentation. Ca cells were considered hyphal if there was any evidence of germ tube formation, regardless of hyphal length.

### Co-aggregation assay

Ca overnight cultures were normalized to an OD600 of 1.0, then diluted 1:5 in a final volume of 1 mL of YPD with 10% FBS in deep-well 96-well plates. “GBS alone” wells were filled with 1 mL YPD + 10% FBS. Where indicated, 5 μg/mL aTC was added to the wells. Each condition was plated in technical quadruplicate. Plates were covered with a gas-permeable plate sealer and incubated shaking at 37°C for 4 hours to induce filamentation. 50 μL of GBS cultures were added to each well, then plates were incubated at room temperature for 2 hours. Plates were centrifuged at 100 g for 6 minutes to pellet the Ca hyphae and any associated GBS, without pelleting non-associated GBS. 25 μL was gently pipetted off the top of each well, serially diluted, and plated for GBS CFU enumeration on THA with 50 μg/mL nystatin. In the “GBS + Ca” wells, this value represents the non-associated GBS. In the “GBS alone” wells, it represents the total amount of GBS in the well, disregarding any rare highly self-aggregating GBS cells. The percentage of GBS associated with Ca was calculated using the following equation:

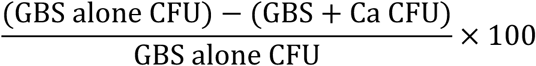

### Biofilm growth

Ca yeast cells were diluted to a concentration of 10^6^ cells/mL and mid-log GFP-GBS at 10^7^ CFU/mL in BHI with 10% FBS in 96-well plates (antibiotic treatment) or coverslip-bottom chamber slides (imaging). Plates were centrifuged at 200 g for 5 minutes, then incubated shaking slowly in a 37°C humidified orbital mixer. After 90 minutes of attachment, culture surfaces were washed with PBS. 200 μL of fresh BHI with 10% FBS was added to each well. Plates were placed back in the shaker and biofilms were allowed to grow for 24 hours (antibiotic treatment) or 48 hours (imaging). For biofilms treated with antibiotics, samples were washed with PBS at 24 hours and fresh media with the indicated concentrations of antibiotic were added to the wells before placing them back in the shaker for 24 hours. Biofilms were then washed with PBS, dissociated and homogenized with vigorous pipetting in 0.25% Triton X-100, and plated for GBS CFU enumeration on THA with 50 μg/mL nystatin. Experiments were performed in technical triplicate. To process samples for imaging, biofilms grown for 48 hours were washed twice with PBS and fixed with 4% paraformaldehyde for 20 minutes. Biofilms were washed three times with PBS, then stained with calcofluor white overnight.

Biofilms were then washed twice with PBS then cleared with the application of SlowFade Glass mounting media. Slides were imaged on a Zeiss LSM780 inverted laser scanning confocal microscope equipped with a 488 nm CW laser and a tunable TPE femtosecond laser.

### Air-liquid interface cell culture

5×10^4^ hVECs were seeded onto 24-well plate transwell membrane inserts with KSFM loaded into the basal compartment. Cells were cultured for two days to allow for attachment and production of a monolayer. Every 48 hours, media was carefully aspirated away from the apical compartment and the media in the basal compartment was changed. Following 4-5 days of culture, media ceased to leak from the basal compartment into the apical compartment, indicating the production and maintenance of tight junctions. hVECs were used for experiments 10-15 days after seeding.

### Adherence assay

All adherence assays were performed in technical triplicate or quadruplicate. hVEC media was changed to fresh KSFM approximately 30 minutes prior to inoculation. For monolayers, 400 μL of KSFM was added to each well and a transwell insert was placed into each well. Ca cultures were normalized to a concentration of 10^6^ cells/mL in KSFM and 100 μL was added to the hVECs into the apical compartment of the transwell insert: directly onto the hVEC cell surface for ALI cultures or physically separated from the hVECs for monolayers. Where indicated, 5 μg/mL aTC was added to the wells.

Conditions without Ca received 100 μL of fresh KSFM. Plates were centrifuged at 200 g for 5 minutes, then incubated at 37°C with 5% CO_2_ for two hours. At the onset of this incubation, GBS overnight cultures, in KSFM for arginine priming experiments and in THB for all other experiments, were diluted 1:10 into fresh media. GBS cultures primed with high arginine were diluted into KSFM with an additional 12 mM arginine added to the media. Once cultures reached mid-log growth, GBS was normalized by OD600 to a concentration of 10^6^ CFU/mL in KSFM. 100 μL was added to the hVECs: into the apical compartment of the transwell for ALI cultures or directly into the well for monolayers, both directly onto the hVEC cell surface. Conditions without GBS received 100 μL of fresh KSFM. Plates were then centrifuged at 200 g for 5 minutes and incubated at 37°C with 5% CO_2_ for 30 minutes. hVECs were washed with PBS five times to remove any non-adherent microbes. For GBS CFU quantification, hVECs were detached with trypsin and dissociated by vigorous pipetting in 0.025% Triton X-100. Homogenized suspensions were diluted and plated on THA with 50 μg/mL nystatin for CFU enumeration. For Ca DNA quantification, transwell membranes were removed and added to screw-cap tubes with Zymo lysis buffer and Zymo BashingBeads. Samples were bead beaten for one minute, five times with ice in between. DNA was isolated using the ZymoBIOMICS DNA Miniprep Kit (Zymo Research, California USA) and quantified by qPCR using primers against the Ca 18S rDNA sequence. A standard curve was created using WT Ca DNA quantified with a Qubit Fluorometer. qPCR plates were loaded in technical triplicate.

### hVEC ALI staining and imaging

Cells were washed with PBS, fixed in 4% paraformaldehyde for 20 minutes, then washed thoroughly with PBS. Cells were then permeabilized in 0.1% Triton X-100 on ice for 15 minutes, washed in PBS, and blocked in 1% BSA for 1 hour. Cells were then stained with phalloidin conjugated to iFluor-555 at a 1:1000 dilution in 1% BSA for 45 minutes and washed with PBS. The membranes were then removed from the plastic transwell inserts and mounted on glass slides using Fluoroshield mounting medium with DAPI. Samples were imaged on a Zeiss LSM780 inverted laser scanning confocal microscope equipped with 488nm, 561nm, and 633nm CW lasers and a tunable TPE femtosecond laser. Images were acquired using hyperspectral imaging with a GaAsP QUASAR detector. Spectral unmixing was performed using ZEN Black software. Image quantification and visualization was performed using Imaris and ImageJ^104^.

### RNA sequencing sample preparation

6-well transwell inserts were seeded with 7×10^5^ hVECs and differentiated for 16 days. Water-tight cell-cell junctions were formed after 15 days of culture. Media was changed in the basolateral compartment and aspirated away from the apical compartment every second day and for the last three consecutive days. For infected hVEC ALI cultures, 2×10^7^ Ca yeast cells (529L) and/or 2×10^8^ GBS CFU at mid-log (COH1) were washed in KSFM and added to the apical compartment in a final volume of 1 mL of KSFM. 1 mL KSFM was added to the apical compartment of uninfected hVEC ALI cultures. For experimental conditions without hVECs, Ca and/or GBS were added to the wells of 6-well plates in a final volume of 1 mL of KSFM. Each experimental condition was performed in technical triplicate. Plates were incubated for 6 hours at 37°C with 5% CO_2_. Following the incubation, supernatants were aspirated off of the hVECs so as only to collect microbes located directly at the hVEC surface. Supernatants from conditions without hVECs were centrifuged to collect microbial pellets which were added back to the wells. All wells were vigorously scraped and pipetted in lysis buffer containing 1% β-mercaptoethanol. The suspension was added to screw-cap tubes with 0.1 mm zirconia beads and bead beaten for 40 seconds twice with ice in between. Tubes were centrifuged and the lysed sample was added to an equal volume of fresh 75% ethanol. Samples were loaded onto Qiagen RNeasy spin columns, washed, and eluted following the manufacturer’s instructions. Eluted RNA samples were treated with Invitrogen DNase following manufacturer’s instructions.

### RNA sequencing

Library preparation was performed using Illumina’s Stranded Total RNA Prep ligation with Ribo-Zero Plus rRNA depletion kit, including custom probes against *C. albicans* rRNAs, and 10-bp unique dual indices. Sequencing was performed on a NovaSeq X Plus with 150×2 reads. BCL convert v4.2.4 was used for demultiplexing, quality control, and adapter trimming. Conditions with GBS alone had ≥6.10×10^6^ read pairs, with Ca and/or hVECs ≥1.71×10^7^ read pairs, and with all three organisms ≥1.59×10^8^ read pairs.

### Transcriptomics data analysis and visualization

STAR^105^ v2.7.10b was used to align reads to the appropriate reference genomes in order of increasing expected RNA abundance in the sample. For example, the GBS+Ca conditions were first aligned to the GBS genome, and the remaining unmapped reads were aligned to the Ca genome. The Ca+hVEC conditions were first aligned to the Ca genome, and the remaining unmapped reads were aligned to the human genome. We used GBS genome NZ_HG939456.1 with sRNAs^106^, Ca Assembly 22^107^, and human genome assembly GRCh38. Samples were filtered based on mapping statistics, which resulted in the removal of three samples from analysis: one Ca+hVEC replicate, one GBS+hVEC replicate, and one GBS+Ca replicate due to a combination of low percentages of mapped reads, high mismatch rates per base, and low average alignment lengths. These three experimental conditions were analyzed in technical duplicate and the other four conditions in technical triplicate. Read counts were generated using featureCounts^108^ v2.0.6, and differential expression and statistics were calculated using EdgeR^109–111^ v3.36.0. Volcano plots were generated with EnhancedVolcano^112^ v1.13.2. GSEA was performed using clusterProfiler^113^ v4.2.2.

### Arginine biosynthesis reporter assay and analysis

Experiments were performed in technical duplicate or triplicate. Ca culture was diluted to 10^5^ cells/mL in KSFM with 40 mM MOPS at pH 7.0 and added to an Ibidi 8-well coverslip bottom chamber slide. For transwell conditions, Ca was added to a MatTek 6-well coverslip bottom plate and a transwell insert was placed in each well. Mid-log GBS was added to the wells of the chamber slide or to the apical compartment of the transwell at 10^6^ CFU/mL. Plates were centrifuged at 200 g for 5 minutes, then incubated at 37°C with 5% CO_2_ for six hours. Wells were imaged on a Keyence BZ-X700 inverted fluorescence microscope equipped with a GFP filter cube (ex 470/40 em 525/50) and a TRITC filter cube (ex 545/25 em 605/70). Images were processed and quantified using ImageJ^104^. The image was thresholded in the yCherry (constitutively expressed) channel to create a selection encompassing the area of the Ca. This selection was transferred to the GFP (expressed under the *ARG1* promoter) channel and the mean fluorescence intensity (MFI) was calculated for GFP.

### Quantitative metabolomics

Amino acids were quantified using ultra performance liquid chromatography on a Waters Acquity UPLC system with an AccQ-Tag Ultra C18 1.7 μm 2.1×100mm column and a photodiode detector array. Analysis was performed using the UPLC AAA H-Class Application Kit (Waters Corporation, Milford MA) according to manufacturer’s instructions. Blanks and standards were run every eight samples, and standards were additionally run at the beginning and end of each session. Results were omitted if the standards deviated by greater than 5%. Mass spectrometry grade chemicals and reagents were used.

### RT-qPCR

2×10^8^ GBS CFU at mid-log (COH1) and, where indicated, 2×10^7^ Ca yeast cells (SC5314) were washed in KSFM and added to the wells of 6-well plates in a final volume of 1 mL of KSFM. Where indicated, wells were supplemented with an additional 12 mM arginine. Each experimental condition was performed in technical triplicate.

Plates were incubated for 6 hours at 37°C with 5% CO_2_. Supernatant from each well was centrifuged to form a pellet. Supernatants were removed and pellets were resuspended in Trizol, added back to the wells, vigorously scraped pipetted, then transferred to screw-cap tubes with 0.1 mm zirconia beads. Samples were bead beaten for 30 seconds five times with ice in between. Lysed and homogenized samples were added to 1/5 volume of chloroform and shaken for 30 seconds to mix. Samples were incubated for 2-3 minutes, then centrifuged at top speed for 15 minutes at 4°C. The aqueous phase was carefully pipetted off and added to a clean tube with 500 μL isopropanol. Samples were incubated for 10 minutes at 4°C, then centrifuged at top speed for 10 minutes at 4°C. Supernatant was discarded and pellets were washed in 1 mL fresh 75% ethanol, then centrifuged at top speed for 5 minutes at 4°C. Pellets were air-dried and resuspended in water at 55°C for 10 minutes. Samples were treated using the TURBO DNase kit following manufacturer’s instructions. RNA was converted to cDNA using the Quanta qScript cDNA synthesis kit following manufacturer’s instructions. Resulting cDNA was used for qPCR with the QuantaBio PerfeCTa qPCR FastMix following manufacturer’s instructions. qPCR plates were loaded in technical duplicate.

