## Supplemental Figures for "A fungal pathobiont promotes *Streptococcus agalactiae* vaginal persistence and pathogenesis through physical and metabolic interactions"

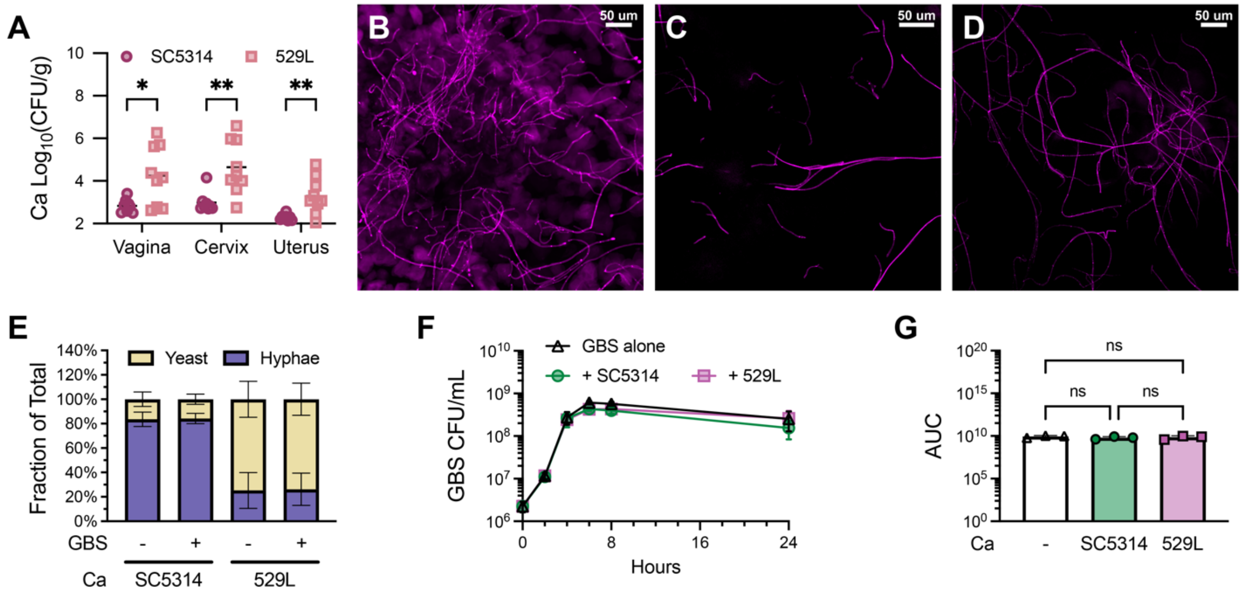


**Figure S1: Ca and GBS outcomes *in vivo* and *in vitro*. (A)** Ca (SC5314 or 529L) burdens in mice co-colonized with GBS (COH1), related to Fig. 1C. n = 8-9 mice per group. **(B)** Vaginal lavage from CD1 mice colonized with Ca (SC5314; magenta) collected one day post inoculation. **(C-D)** Vaginal lavage from C56BL/6 mice colonized with Ca strain SC5314 (C) or 529L (D) was collected seven days post inoculation and stained with calcofluor white. **(E)** Ca (SC5314 or 529L) was grown in KSFM alone or with GBS (COH1) for one hour and imaged. The percentage of cells undergoing filamentation was calculated. Mean ± SEM from three independent experiments. 30 images per group. **(F-G)** GBS (COH1) was grown in KSFM alone or in co-culture with Ca (SC5314 or 529L) and plated for GBS CFU enumeration. CFUs over time (F) and area under the curve (G) are shown. Mean ± SEM from three independent experiments. 9 technical replicates. p values determined by t test (A), two-way ANOVA (E), or one-way ANOVA (G) with Holm-Sidak's multiple comparisons test. * $p<0.05$; ** $p \leq 0.01$

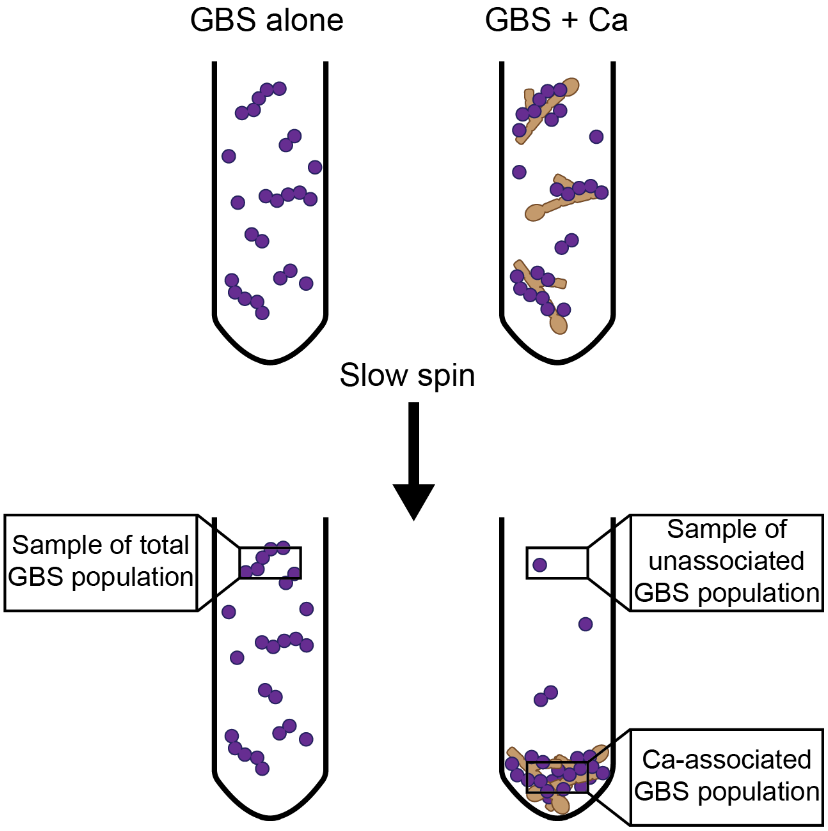


$$\% \text{ Co-Aggregation} = \frac{\text{Total - Unassociated}}{\text{Total}} \times 100 = \frac{\text{Associated}}{\text{Total}} \times 100$$

**Figure S2: Schematic of co-aggregation assay.** Equal quantities of GBS are inoculated into wells with and without Ca. Samples are centrifuged at a low speed such that the GBS is minimally disturbed and does not form a pellet, but maximal Ca flocculates and forms a pellet. We assume that the GBS is homogenously distributed throughout the well in the “GBS alone” condition and that any non-associated GBS is homogenously distributed in the supernatant of the “GBS + Ca” condition. A small volume is pipetted off the top of each well and plated for GBS CFU enumeration. These values are used to calculate the total amount of GBS in the well, extrapolated from the “GBS alone” condition, and the total amount of non-associated GBS in the well, extrapolated from the “GBS + Ca” condition. These values are then used to calculate the percentage of the total GBS population that is physically associated with Ca. This method normalizes for varying rates of self-aggregation in the event that some isolates may form aggregates that are sufficiently dense to flocculate under these conditions. This assay calculates any enhancement in GBS aggregation attributed to physical interactions with Ca.

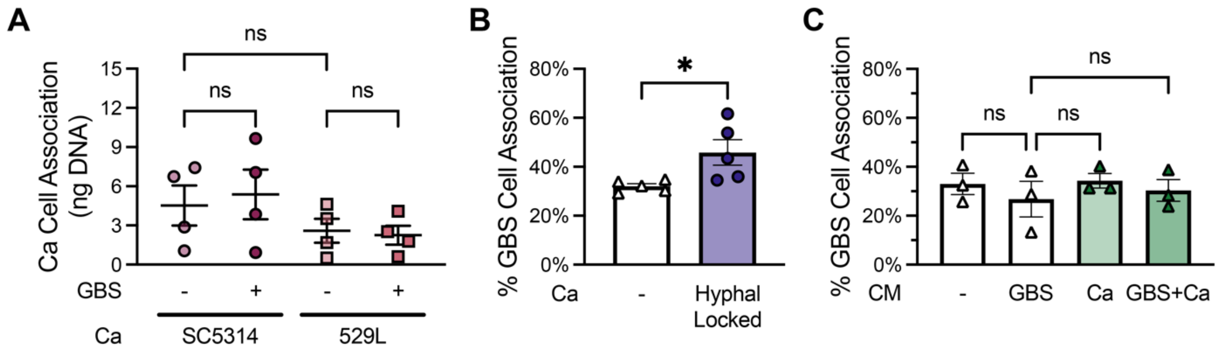


**Fig. S3: Increased GBS adherence is contact-dependent. (A)** Adherent Ca (SC5314 or 529L) quantified by qPCR for Ca 18S rRNA. Mean ± SEM from four independent experiments. 12 technical replicates. **(B)** Filamentation was induced in Ca (TetO-*NRG1*) by the addition of 5 ug/mL aTC. GBS (COH1) was inoculated onto hVEC ALI cultures alone or with Ca hyphae and incubated for 30 minutes. Cultures were washed, dissociated, and plated for GBS CFU enumeration. Mean ± SEM from five independent experiments. 15 technical replicates per group. **(C)** GBS (COH1) was inoculated onto hVEC ALI cultures in KSFM or KSFM conditioned with GBS (COH1) or Ca (SC5314) monocultures or a co-culture. Cultures were washed, dissociated, and plated for GBS CFU enumeration. Mean ± SEM from three independent experiments. 9 technical replicates per group. p values determined by one-way ANOVA with Holm-Sidak's multiple comparisons test (A, C) or t test (B). * $p<0.05$

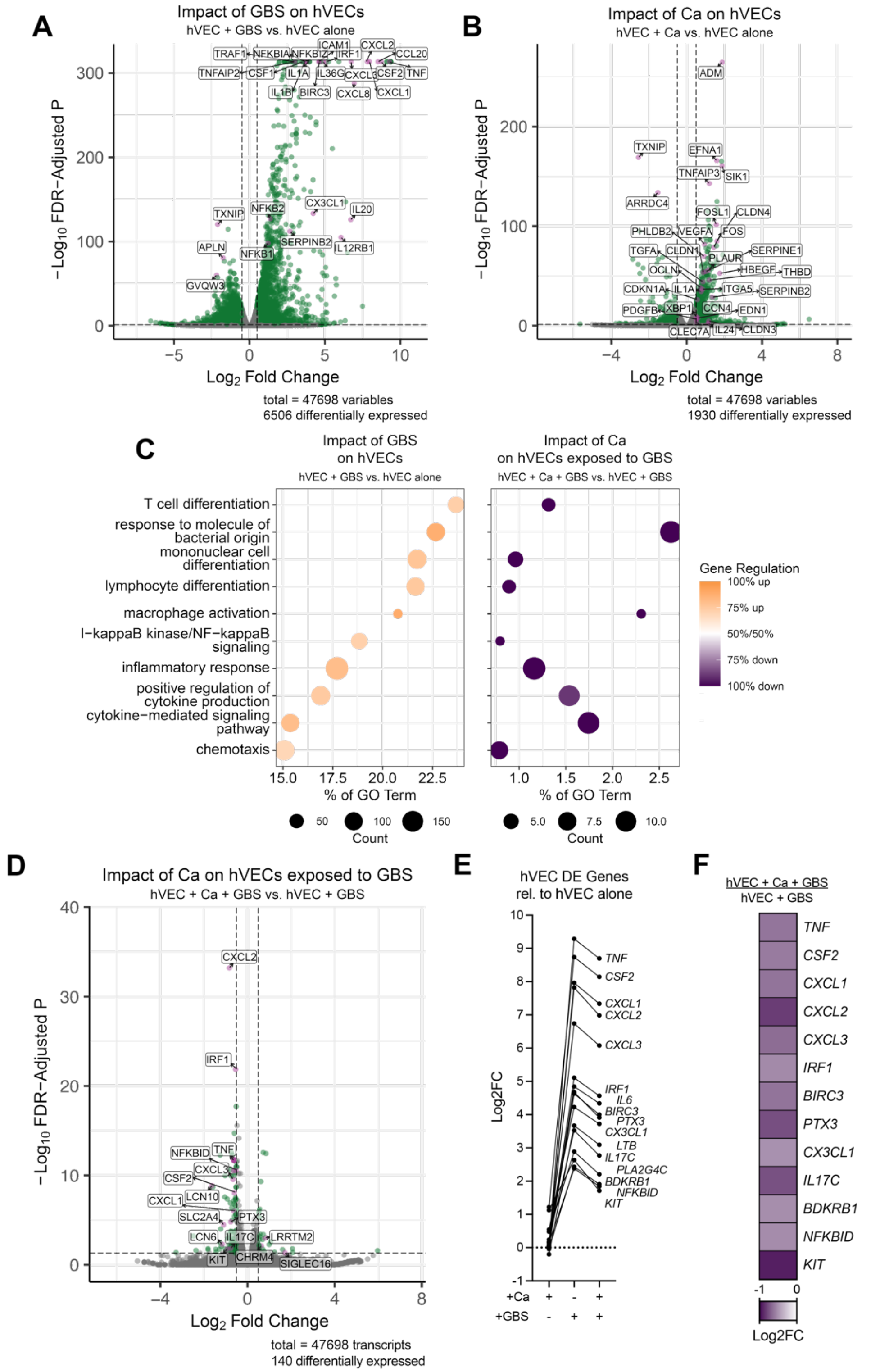


**Fig. S4: Host transcriptional response to GBS and Ca. (A)** hVEC DE genes with GBS compared to hVEC alone. **(B)** hVEC DE genes with Ca compared to hVEC alone. **(C)** hVEC DE genes were analyzed for enrichment of GO terms. Dot size represents the number of DE genes within a GO term. X-axis represents the proportion of all genes within a GO term that are DE. Dot color represents the proportion of DE genes within a GO term that are up or downregulated. **(D)** hVEC DE genes with Ca and GBS compared to with GBS only. **(E)** Log2(Fold Change) of select hVEC genes with Ca, GBS, or Ca and GBS together compared to hVEC alone. **(F)** hVEC DE genes with Ca and GBS compared to with GBS only.

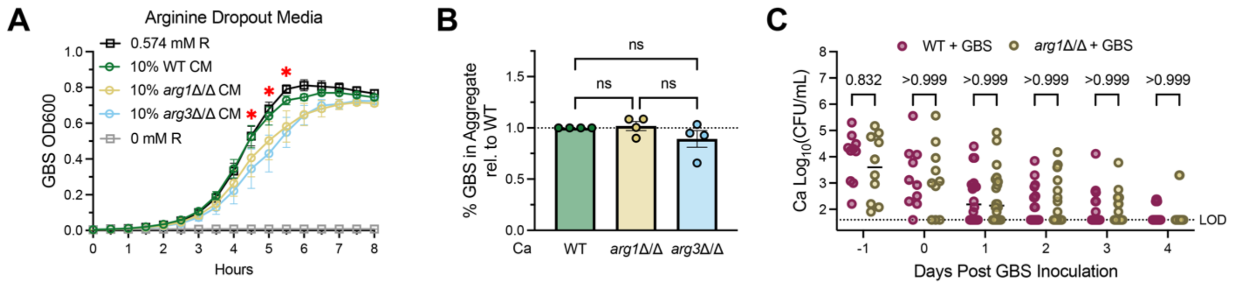


**Fig. S5: Arginine biosynthesis does not impact co-aggregation or vaginal colonization. (A)** GBS (COH1) was grown in chemically defined medium (CDM) without arginine. CDM was supplemented with Ca (SC5314 WT, *arg1*Δ/Δ, or *arg3*Δ/Δ) conditioned KSFM, untreated KSFM, or 0.574 mM arginine. Asterisks indicate p<0.05 between 10% WT CM and 10% *arg1*Δ/Δ or *arg3*Δ/Δ CM. Mean ± SEM from five independent experiments. 15 technical replicates per group. **(B)** GBS (COH1) co-aggregation with Ca (SC5314 WT, *arg1*Δ/Δ, or *arg3*Δ/Δ). Mean ± SEM from four independent experiments. 16 technical replicates per group. **(C)** Ca (SC5314 WT or *arg1*Δ/Δ) burdens in the vaginal lavage of mice co-colonized with GBS (COH1), related to Fig. 6K. n = 17-19 mice per group. p values determined by two-way repeated measures ANOVA (A), one-way ANOVA (B), or two-way ANOVA (C) with Holm-Sidak's multiple comparisons test. * p<0.05
